# Pathway-centric multi-omics and functional precision medicine reveal shared drug vulnerabilities in heterogeneous adult Wilms tumor

**DOI:** 10.64898/2026.06.23.734042

**Authors:** Minttu Polso, Romika Kumari, Tamara J. Luck, Piia Mikkonen, Katja Välimäki, Ruusu-Maaria Merivirta, Minna Malmstedt, Johanna Lehtonen, Erika Romppanen, Sara Kuusela, Antti Hassinen, Janna Saarela, Teijo Pellinen, Panu M. Jaakkola, Pia Suonpää, Petrus Järvinen, Olli Kallioniemi, Tuomas Mirtti, Antti Rannikko, Vilja M. Pietiäinen

**Affiliations:** Institute for Molecular Medicine Finland-FIMM, Helsinki Institute for Life Sciences-HiLIFE, University of Helsinki, Finland; iCAN Flagship, University of Helsinki, Finland; Department of Oncology, HUS Helsinki University Hospital, Helsinki, Finland; Department of Medical Genetics, Oslo University Hospital, Oslo, Norway; Norwegian Centre for Molecular Biosciences and Medicine-NCMBM, Oslo, Norway; Science for Life Laboratory and Department of Oncology and Pathology, Karolinska Institutet, Stockholm, Sweden; Department of Pathology, HUS Helsinki University Hospital, Helsinki, Finland; ONCOSYS Research Program of Systemic Oncology, Faculty of Medicine, University of Helsinki, Finland; Turku University and Turku University Hospital, Turku, Finland; Finnish Cancer Institute, Helsinki, Finland

**Keywords:** Wilms tumor, nephroblastoma, functional precision medicine, drug testing, multi-omics, intratumoral heterogeneity

## Abstract

Wilms tumor, *i.e.,* nephroblastoma, is rare in adults and lacks standardized treatment, complicating clinical decision-making. Within the functional precision medicine study (DEDUCER), we profiled two spatially distinct tumor regions (T1 and T2) of an adult Wilms tumor patient using integrated histopathology, whole-exome sequencing, FFPE transcriptomics, and *ex vivo* drug screening of short-term cultured patient-derived cancer cells (PDCs) with 528 compounds. Genomic profiling revealed a truncal ASXL1 frameshift and shared *F7, UBA1,* COL21A1, and ATM variants alongside region-specific alterations: a TP53 mutation and broad copy-number (CN) gains in T1, versus ARID1A and KMT2D stop-gains in copy-neutral T2. Transcriptomics of tumor areas identified convergent activation of the G2/M checkpoint, E2F targets, and mitotic spindle programs across regions, consistent with high proliferation and partially comparable biomarker signatures to those observed in an open-source pediatric Wilms tumor dataset (n = 130). Functional assays uncovered distinct and shared drug vulnerabilities: although ATM alterations were present in both tumors, T1 PDCs showed selective sensitivity to topoisomerase I and BCL-2 inhibition in the context of an additional T1-specific TP53 alteration, while broader single-agent sensitivity and stronger drug synergies were observed in T2.

Pathway-centric data integration indicated that differential gene expression and copy-number gains, rather than single mutations alone, better predicted ex vivo drug responses, revealing actionable shared dependencies despite pronounced spatial heterogeneity and establishing a translational framework for individualized management in this rare disease.

**HIGHLIGHTS:**

– In the adult Wilms tumor, multi-region genomics revealed a truncal ASXL1 frameshift together with F7, UBA1, COL21A1 and ATM mutations across two tumor regions (T1 and T2), as well as region-specific alterations: TP53 mutation and widespread copy-number gains in T1, versus ARID1A and KMT2D stop-gains in copy-neutral T2, illustrating spatial heterogeneity.
– Transcriptomics showed convergent activation of E2F targets, G2/M checkpoint, and mitotic spindle programs in both regions, consistent with high proliferation and aligning with Wilms tumor signatures (TARGET dataset); these pathways were associated with higher *ex vivo* drug sensitivity scores.
– Functional drug sensitivity testing of patient –derived cancer cells *ex vivo* uncovered distinct and shared vulnerabilities: Although both tumors shared an ATM mutation, T1-specific TP53 alteration and death-pathway/stress-response alterations may underlie selective sensitivity to topoisomerase I inhibitors and BCL-2 inhibition.
– Clinically relevant combinations, including vincristine plus dactinomycin and doxorubicin plus dactinomycin, showed ex vivo synergy. These findings are consistent with the patient’s more than five-year relapse-free outcome following vincristine, doxorubicin, and dactinomycin treatment combined with surgery, supporting the translational relevance of the ex vivo drug testing approach.
– Pathway-centric integration (copy-number gains and differential expression) predicted drug response better than single-gene biomarkers. Overall, pathway-level dependencies provide robust, actionable targets despite genomic and phenotypic heterogeneity in adult Wilms tumor.

## INTRODUCTION

Wilms tumor (WT, nephroblastoma) is a rare disease in adults, with an annual incidence of only 0.2 cases per million (Mitry et al. 2006). The tumor is often characterized by chromosomal alterations on chromosome 11p13, including the Wilms tumor suppressor gene, WT1, which is a zinc-finger DNA-binding transcription factor expressed throughout kidney development (Pritchard-Jones et al., 1990, Larsson et al. 1995, Ozdemir and Hoehenstein, 2014). Loss of heterozygosity (LOH) at chromosomes 1p and 16q, gain of chromosome 1q, and high telomerase (TERT) expression are associated with adverse prognosis and inferior survival in favorable-histology Wilms tumor (Grundy et al., 2005; Park et al., 2020; Jablonowski et al., 2022). In adults, Wilms tumor is typically diagnosed at more advanced stages and lacks standardized treatment guidelines, resulting in reliance on pediatric protocols given limited adult-specific molecular and therapeutic data. The TARGET Wilms tumor project represents the most comprehensive molecular characterization effort to date for high-risk pediatric Wilms tumor, identifying recurrent genetic alterations and dysregulated developmental pathways at the cohort level (Gadd et al., 2017). Through integrated multi-omics analyses, TARGET revealed that diverse genetic alterations converge on key developmental and cell-cycle–associated pathways, highlighting pathway-level dysregulation despite substantial inter-tumor genetic heterogeneity. However, while TARGET has provided critical insights into the genomic and transcriptomic landscape of Wilms tumor, it was not designed to resolve functional drug dependencies or to assess spatial intratumor heterogeneity within individual tumors. The project identified recurrent mutations and alterations in genes involved in early kidney development, including WT1, SIX1, SIX2, MYCN, and genes involved in microRNA processing (dbGaP sub-study ID: phs000471).

In a comprehensive genomic and multi-omics analysis of 117 pediatric Wilms tumors, additional recurrent mutations were discovered in genes not previously implicated in Wilms tumor pathogenesis, such as BCOR, BCORL1, NONO, MAX, COL6A3, ASXL1, MAP3K4, and ARID1A (Gadd et al., 2017). Copy number alterations were also identified, including 1q gain, MYCN amplification, LIN28B gain, and MIRLET7A loss. A separate study published in 2022 analyzed 14 adult Wilms tumors (patients aged 17–46 years) using expanded targeted sequencing. This study identified clinically actionable mutations such as BRAF V600E, as well as alterations commonly seen in pediatric Wilms tumor, including mutations in WT1, ASXL1, NSD2, and loss of 11p (Argani et al., 2022).

Although adult and pediatric Wilms tumors share similar histological features, adults often present with more advanced disease and experience poorer outcomes, largely due to delayed recognition and the absence of standardized treatment protocols (Saltzman et al., 2020). Initiatives have already been undertaken for pediatric Wilms tumor to integrate multi-omics data into clinical decision-making, paving the way for more individualized treatment pathways. However, such efforts remain limited in adult Wilms tumor due to the rarity of the disease. As there are no established treatment guidelines for adults, patients are treated on an individual basis, primarily by following the treatment protocols for pediatric Wilms tumor (Modi et al., 2016). These regimens typically involve multimodal therapy combining surgery, radiotherapy, and multi-agent chemotherapy, most commonly built on vincristine–dactinomycin backbones with the addition of doxorubicin, platinum compounds, or alkylating agents in higher-risk disease. However, adult patients experience higher rates of treatment-related toxicity compared with children, often requiring dose reductions or treatment modifications. Moreover, the efficacy and tolerability of these pediatric-derived regimens have not been systematically evaluated in adults, as prospective or randomized studies are not feasible for this rare and biologically heterogeneous disease (Sakthivel et al., 2023).

Based on recent studies, the patient-derived cancer cells (PDCs) provide unique resources for testing targeted therapies and can also be further utilized to identify and validate novel treatment strategies aimed at improving patient outcomes. For example, in pediatric Wilms tumor models, PDCs, particularly from MYCN-driven tumors, have been shown to exhibit specific drug sensitivities (Götz et al., 2025). Another study established a library of 45 patient-derived xenograft (PDX) models from Wilms tumor samples, demonstrating that these models reliably reproduced histologic, molecular, and chemotherapy response features (Murphy et al., 2019).

This study aimed to characterize the intratumoral molecular profile and analyze potentially shared drug sensitivities between two spatially distinct regions (T1 and T2) of an adult Wilms tumor using multi-omics approaches and *ex vivo* PDC models in functional drug testing (see **Fig. 1A**). To enhance generalizability beyond a single case, we benchmarked our transcriptomic findings against the TARGET Wilms Tumor dataset (n = 130 pediatric cases), observing concordant enrichment of cell-cycle programs (G2/M checkpoint, E2F targets and mitotic spindle) and overlapping gene signatures (e.g., *SIX2*, *BUB1B*, *COL2A1*). The integrative analysis revealed spatial genomic heterogeneity with convergent pathway activation and identified shared and region-specific drug sensitivities and synergies. By integrating multi-omics with *ex vivo* pharmacologic profiling and cross-referencing gene expression against the TARGET cohort, this work advances understanding of adult Wilms tumor biology and identifies clinically relevant, targetable pathways. These insights enable more precise therapeutic prioritization for individualized care in this heterogeneous and rare disease.

**Figure 1.**
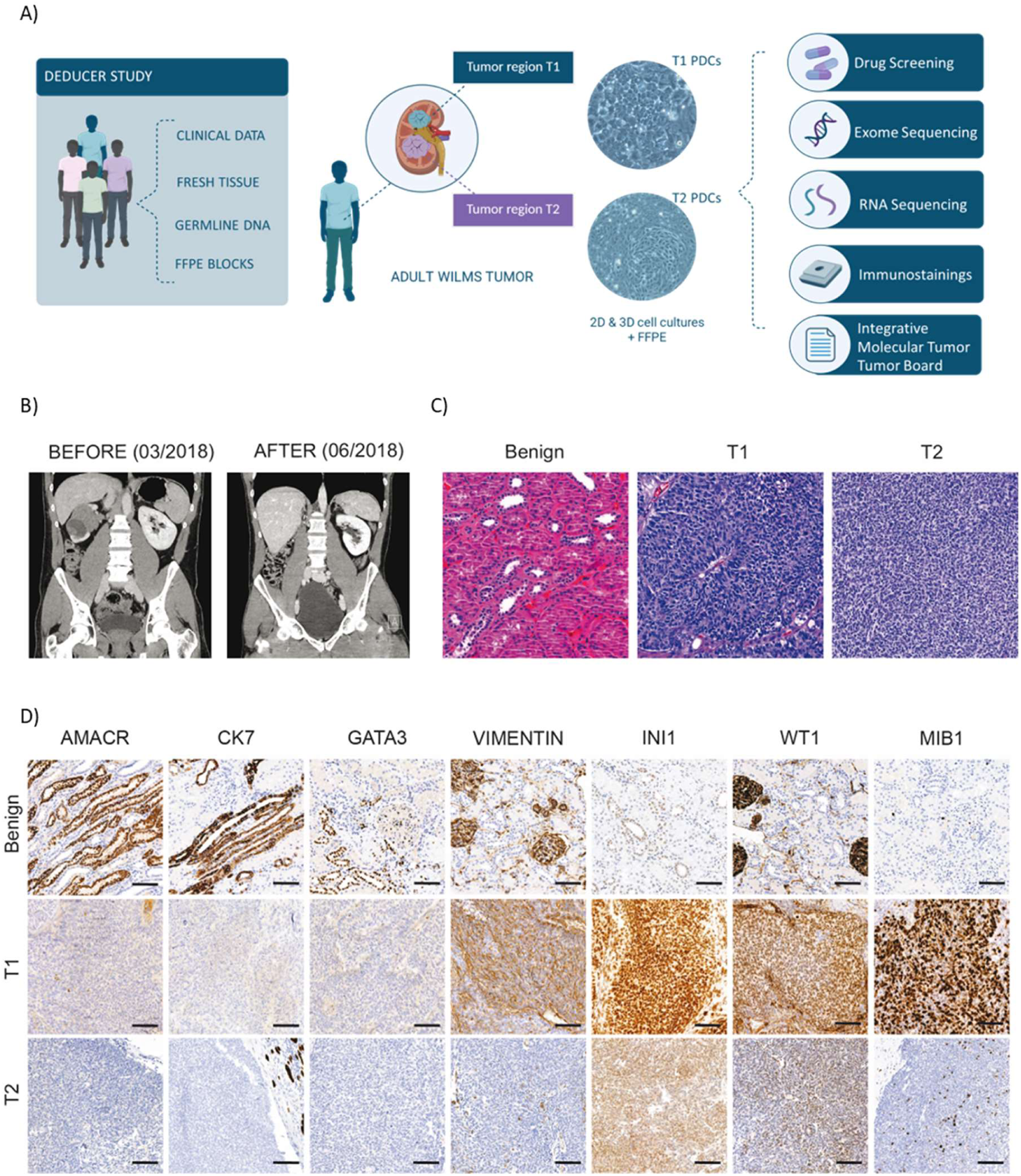
Characterization of tumor regions T1 and T2 in an adult Wilms tumor case. **A)** Overview of the study design within the DEDUCER study, with a subcohort of renal cancers, ‘n-of-1’ clinical trial, NCT02994758), incorporating clinical data, fresh tumor/benign tissue, germline DNA, and formalin-fixed paraffin-embedded (FFPE) tissue samples for functional precision medicine in real-time. T1 and T2 tumor samples were used to establish 2D and 3D patient-derived cancer cell (PDC) models. Exome sequencing was performed on both the original tissue and the derived PDCs, while FFPE tissue was used for immunohistochemistry (IHC) and transcriptomic profiling. **B)** Pre– and postoperative CT scans, showing the nephrectomy of the kidney with tumor mass. **C)** Histopathological examination with hematoxylin and eosin (H&E) staining of benign kidney tissue, T1 and T2, highlighting differences in tissue architecture. Local anaplasia was observed in the T1 section. **D)** Immunohistochemical staining of benign, T1 and T2 tumor tissues for AMACR, CK7, GATA3, Vimentin, INI1, WT1, and MIB1, illustrating distinct biomarker staining patterns between the two tumor regions (see also Supp. Fig. 1 for additional profiling). Scale bars 100 μm. (BioRender.com was used to create Fig. 1A).

## RESULTS

### Pathological biomarker profile confirms Wilms tumor

The patient (an adult male aged between 30–40 years) with localized renal tumor was enrolled in the DEDUCER study (NCT02994758) and underwent radical nephrectomy at Helsinki University Hospital for a right renal mass (5.5 × 4.0 cm) **(Fig. 1B)**. The surgical specimen was dissected routinely, and samples from macroscopically separate tumor regions T1 and T2, benign kidney, and hilar structures were collected for diagnostics and research. No lymphadenectomy was performed, and no lymph nodes were identified in the perirenal fat.

To investigate intratumoral heterogeneity and potential therapeutic vulnerabilities, two histologically distinct tumor regions (T1 and T2) were sampled at the time of surgery for whole-exome sequencing, RNA sequencing, and *ex vivo* PDC drug sensitivity and resistance testing (DSRT) (**Fig. 1A**), as well as examined for pathological biomarkers of Wilms tumor.

Routine histopathology evaluation confirmed a diagnosis of Wilms tumor (SIOP nephroblastoma stage II) with expression of Wilms tumor-specific markers **(Fig. 1C-D).** Histological subtypes included blastemal, epithelial, and stromal components; however, the immunohistochemistry revealed differences between the two tumor regions. T1 showed strong expression INI1/SMARCB1 (SWI/SNF chromatin remodeler), PAX8 (paired box transcription factor of renal lineage), and TP53 (tumor suppressor), together with focal WT1 (Wilms tumor 1, zinc-finger transcription factor) positivity and clear vimentin (mesenchymal marker) positivity in blastemal and epithelial areas, accompanied by a high proliferative index (MIB-1/Ki-67: 50–70%). In contrast, T2 demonstrated overall weaker marker expression and a more homogeneous blastemal profile (**Fig. 1D**). All samples were negative for CK7 (cytokeratin 7), GATA3 (GATA binding protein 3), and AMACR (alpha-methylacyl-CoA racemase), supporting the diagnosis of Wilms tumor and arguing against alternative renal tumor entity. No pathogenic germline variants in Wilms tumor–associated genes were identified in clinical panel sequencing.

To further characterize tumor heterogeneity, extended immunohistochemical profiling was performed (**Supp. Fig. 1**). Both regions expressed CD56/NCAM1 (neural cell adhesion molecule) and SIX2 (sine oculis homeobox 2; nephron progenitor transcription factor), with stronger SIX2 staining in T2, while CD133/PROM1 (stem/progenitor marker) was more prominent in T1. In contrast, CDKN1A/p21 (cell-cycle inhibitor), SOX2 (SRY-box transcription factor), and ALDH1 (aldehyde dehydrogenase 1; stemness marker) were negative in both regions.

### Cancer tissue regions harbor both known and novel mutations and copy number alterations

Based on the observed histological and immunophenotypic differences, both tumor tissues and matched T1 and T2 PDCs were subjected to integrated genomic and functional analyses. Somatic mutations and copy number alterations (CNAs) of tumor samples were analyzed using exome sequencing data. Notably, T1 exhibited a higher mutational burden than T2 (1.18 vs 0.62 mutations/Mb).

Both shared and tumor region-specific genetic alterations were observed between T1 and T2. Among the earlier reported mutations in TARGET Wilms tumor (WT) dataset (Cerami et al., 2012), F7 (p.Arg304Trp) and ASXL1 (p.Thr600ProfsTer103) were shared by both T1 and T2, whereas TP53 (p.Gln136Glu, VAF 0.981) and FAM83B (p.Arg910LysfsTer49) were unique to T1, and KMT2D (p.Gln211Ter) and ARID1A (p.Tyr154Ter) were present in T2 only **(Fig. 2A)**. TP53 is considered a likely pathogenic mutation with its known frequency of ∼31% in Wilms tumors (TARGET-WT). The TP53 mutation identified in T1 was associated with strong nuclear TP53 staining in the corresponding tumor tissue **(Fig. 2B)**, and a similar pattern was also observed in the matching T1 PDCs (**Fig. 2C**). The higher TP53 protein expression in T1 PDCs compared to T2 was further shown by Western Blotting, as described below PDCs **(Fig. D)**.

**Figure 2.**
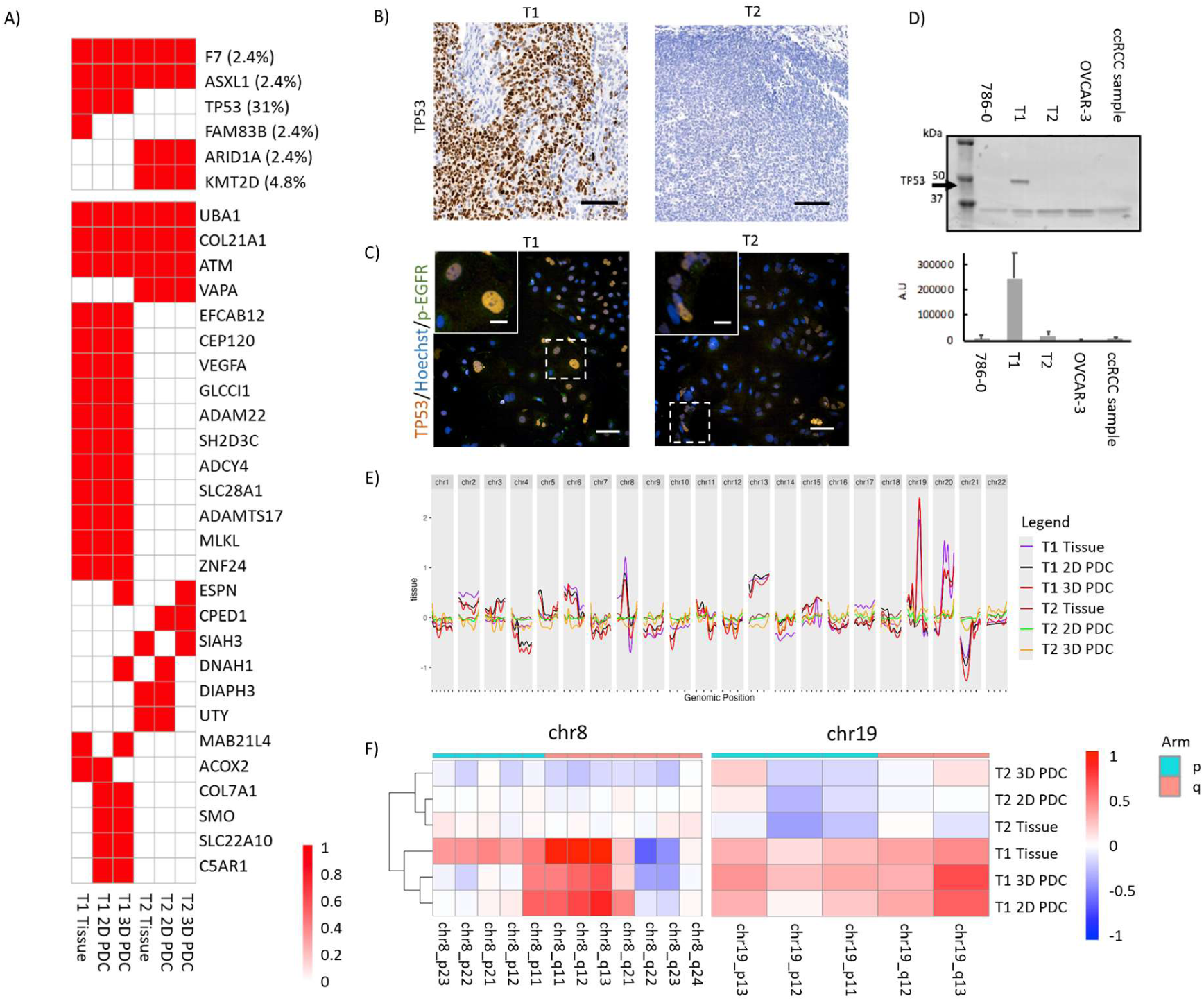
Molecular characterization of patient tumor tissues and patient-derived *ex vivo* cell cultures from tumor regions T1 and T2. A) Mutation status of selected genes including novel mutations and those overlapping with TARGET-WT that were mutated in ≥2 samples across tumor samples, highlighting key alterations (red denotes the presence of a mutation). The frequency of mutations in the TARGET-WT dataset is indicated in parentheses for genes overlapping with TARGET-WT. **B)** IHC-staining for TP53 in tumor tissues from T1 and T2. Scale bars: 100 µm **C)** Immunofluorescence staining of TP53 (orange) together with p-EGFR (green) and Hoechst (blue) in patient-derived cell cultures from T1 and T2, demonstrating differential TP53 protein expression in T1, consistent with the TP53 mutation detected in T1. Scale bars: 100 µm (overview), 25 µm (zoom-ins). **D)** Western blot analysis of TP53 expression in T1 and T2 PDCs, with human renal cell carcinoma cell line 786-O, human ovarian cancer cell line OVCAR-3, and a clear cell renal carcinoma (ccRCC) reference sample from the DEDUCER cohort included as controls. Corresponding densitometric quantification is shown below. TP53 was highly expressed in T1 PDCs compared to the other samples. E) Genome-wide CNA plots showing relative amplifications and deletions across chromosomes in different sample types. F) Cytoband-level CNA heatmaps for chromosomes 8 and 19, highlighting recurrent amplifications and deletions identified in the different sample types.

Additionally, several novel mutations, such as UBA1, COL21A1, and ATM were present in both T1 and T2, along with some unique mutations in either part of the tumor **(Fig. 2A)**. Interestingly, the ATM missense mutation shared between T1 and T2 showed a markedly higher VAF in T1 (p.Leu2890Ile, VAF 0.89) than in T2 (VAF 0.29). In addition, T2 harbored a second ATM mutation that was absent from T1 (p.Ile2888Thr, VAF 0.50). Together, these findings indicate differential mutational patterns affecting ATM between T1 and T2 despite involvement of the same gene. **(Supp. Fig. 2)**.

The somatic CNA analysis uncovered that T1 exhibited amplifications on chromosomes 2, 6, 8(p), 13, 19, and 20(q), and deletions on 8(q), 10, and 21(p) (**Fig. 2E-F)**. Notably, amplifications of 20(q) have previously been linked to renal cancer in the TracerX study (Turajlic et al., 2018). A prominent gain at chr 19q13.33, encompassing the Wilms tumor predisposition gene *FWT2* (McDonald et al., 1998), was identified in T1 tumor tissue (**Fig. 2E-F)**. T2 had a copy-number – neutral profile; no broad deletions or amplifications were detected.

To determine the functional and clinical relevance of the identified genomic alterations, we generated integrative Molecular Tumor Board (iMTB) reports for each sample. Interestingly, only one out of the four functionally relevant and predicted oncogenic mutations identified (as per the standard operating procedures from ClinGen/CGC/VICC (Horak et al., 2022)) were shared between T1 and T2: In T1, the *ASXL1* frameshift and *TP53* missense mutations were predicted oncogenic, while in T2, besides the shared *ASXL1* mutation, *ARID1A* and *KMT2D* stop-gain variants were identified (**Table 1**). Potential clinical significance as a predictive biomarker was attributed only to the *TP53* variant, which has been linked to sensitivity to pazopanib and vorinostat (in solid tumors), VEGF/VEGFR inhibitors (any cancer), chemotherapy (gastric adenocarcinoma and non-small cell lung cancer), and alemtuzumab (leukemia), as well as resistance to tamoxifen (breast cancer) and MDM2-inhibitor RG7112 (leukemia). In addition, several clinically relevant CNAs were identified in T1, including the MET gain, which was attributed to strong clinical significance, as it has been linked to MET inhibitor sensitivity in other kidney cancers. The CCND1 gain was classified as Tier 2, indicating potential clinical relevance based on its established role in cell-cycle regulation. In addition, potential clinical significance was attributed to 16 other gene amplifications in T1 (PIK3CA, ERBB2, VEGFA, TERT, FGFR1, FGF3, FGF4, TOP2A, TOP1, CCND1, CCNE1, MYCN, REL, IGF2, NCOA3, RSF1; (**Supp. Table 1**). Ten of these had again previously been identified as predictive biomarkers, linked to the response of 35 different drugs or sets of drugs in cancer; eight were prognostic biomarkers linked to poor outcome (**Supp. Table 1**).

**Table 1.**
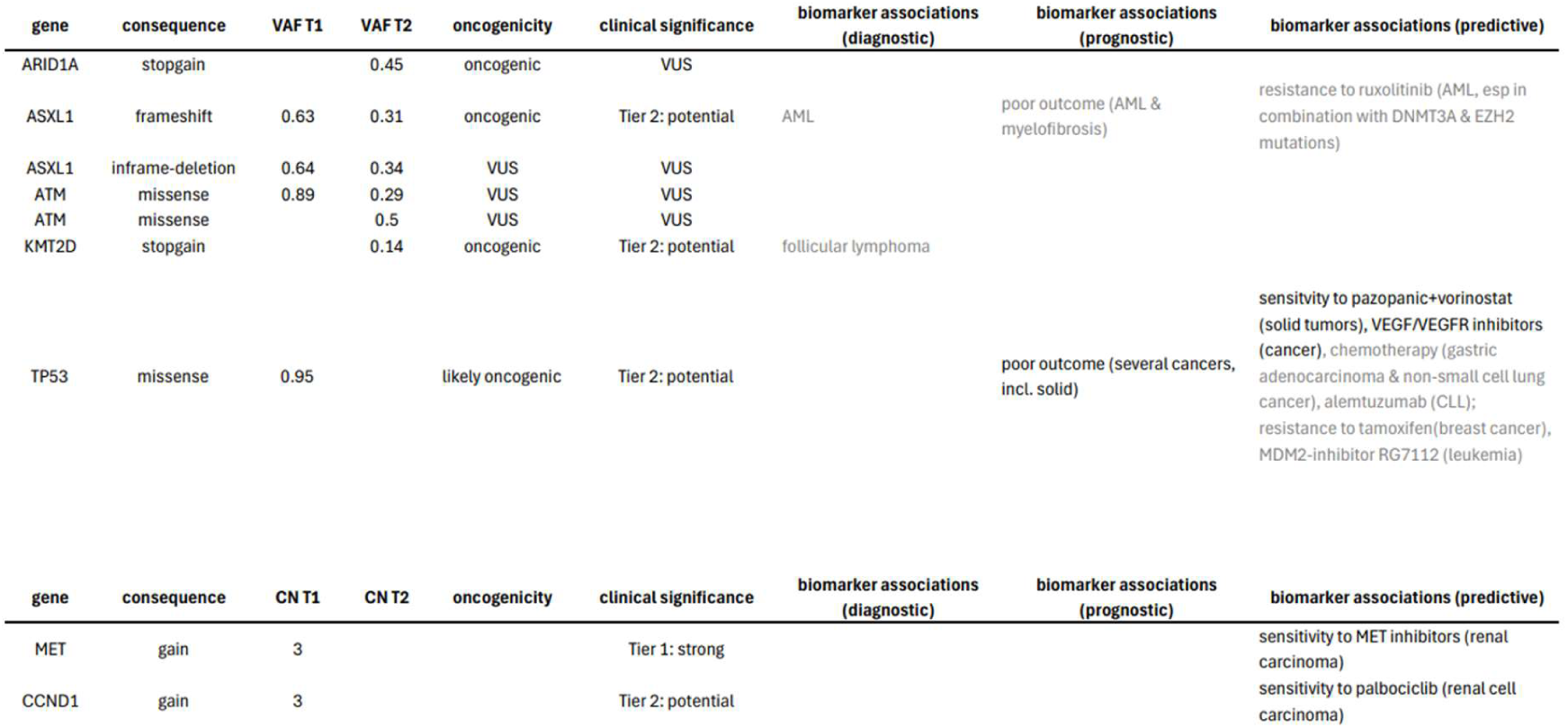
Summary of somatic SNVs, indels, and copy-number alterations classified as oncogenic or associated with Tier 1–2 biomarkers in T1 and T2. As T1 contained Tier 2 CNAs associated with more than 80 biomarkers, only those associated with cancer/solid cancer/kidney are listed below. The complete list can be found in **Supp. Table 1.** For SNVs/Insertions/Deletions, all biomarkers are listed, but those associated with anything other than cancer/solid cancer/kidney are grayed out.

### Transcriptomic profiling shows enrichment in G2M checkpoint and E2F targets for both tumor regions

To characterize differences between tumorigenic (T1 and T2) and benign kidney regions at the transcriptomic level, we performed HTG EdgeSeq profiling with the HTG Transcriptome Panel on tumor cells obtained from FFPE samples, where FFPE samples from regions of interest (T1, T2, and benign tissue) were collected using three methods: (1) scalpel scraping directly from the tissue slide, (2) laser microdissection without paraffin removal, and (3) laser microdissection after paraffin removal and tissue staining (**Fig. 3A**). The correlation analysis of the transcriptomic profiles obtained from these three different methods revealed significantly strong reproducibility of the gene expressions profiles with Pearson correlation coefficients varying from 0.87-0.98, which was further supported by principal component analysis (PCA) **(Supp. Fig. 3 A-B)**. Subsequently, these samples were used as technical/methodological replicates for differential gene expression analysis.

**Figure 3.**
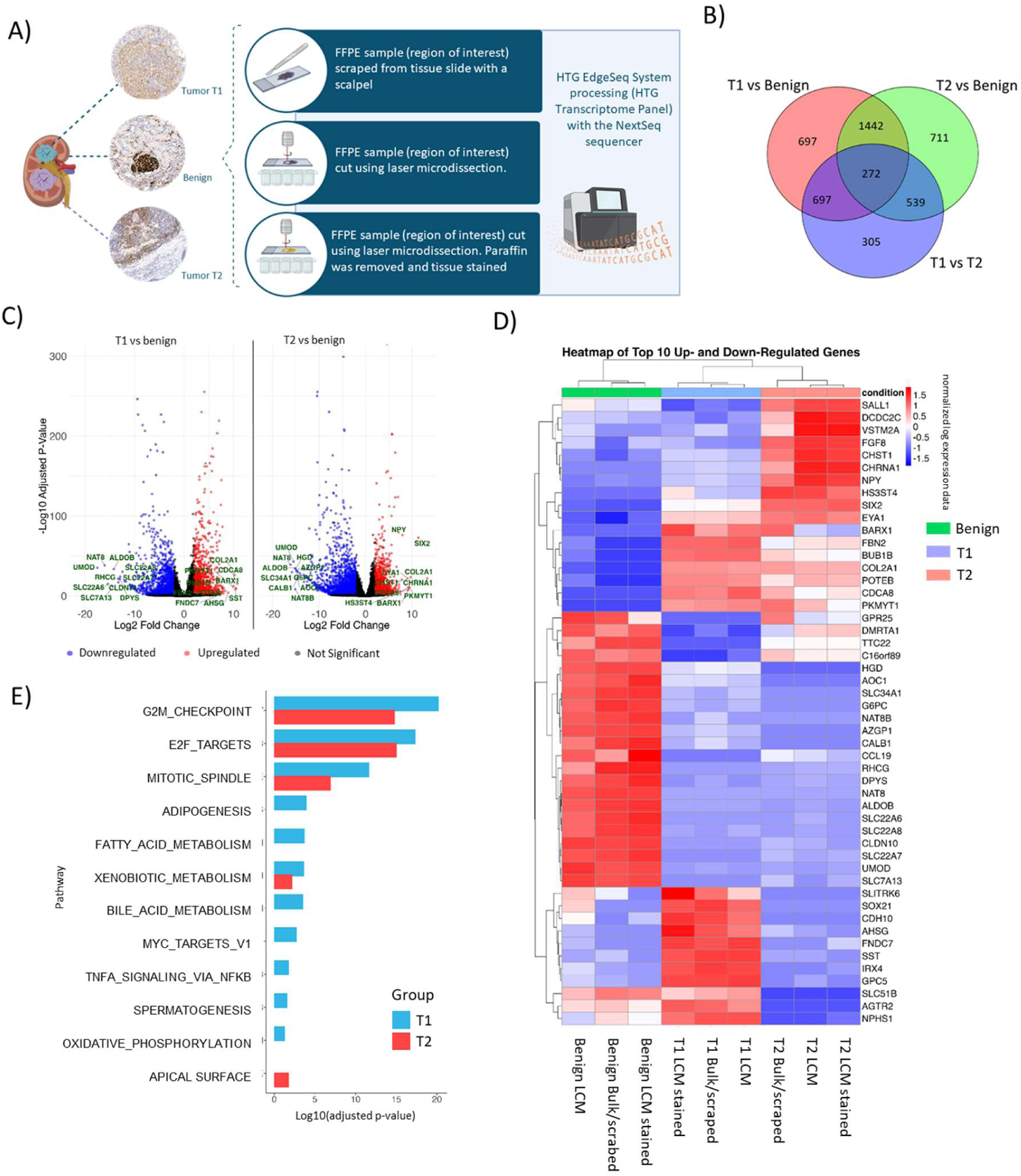
Transcriptomic profiling of FFPE samples from tumor regions (T1, T2) and benign kidney tissue. A) FFPE samples from regions of interest (T1, T2, and benign tissue) were collected using three methods: (1) scalpel scraping directly from the tissue slide, (2) laser microdissection without paraffin removal, and (3) laser microdissection after paraffin removal and tissue staining. **B)** Venn diagram showing overlap of differentially expressed genes between comparisons: T1 vs. benign (blue), T2 vs. benign (red), and T1 vs. T2 (green). **C)** Volcano plots illustrating significantly differentially expressed genes in T1 vs. benign (left) and T2 vs. benign (right), highlighting the top 10 up and down DEG genes in green. **D)** Heatmap of the top 10 upregulated and downregulated genes <u>(filtered using log₂(FC))</u> across all conditions (T1, T2, benign). **E)** Gene set enrichment analysis (Hallmark pathways) of differentially expressed genes in T1 (blue) and T2 (red) compared to benign tissue.

The differential gene expression analysis revealed substantial transcriptional changes across tumor regions (T1 and T2) compared to the benign tissue. A total of 2740 and 2964 genes were significantly differentially expressed (adjusted p-value (Benjamini–Hochberg FDR) < 0.01 and |log₂(FC)| > 2) in the T1 and T2 tumor regions versus benign tissue, respectively. Overall, the analysis revealed 1442 genes differentially regulated in both T1 and T2 relative to benign tissue, indicating shared tumor-associated transcriptional changes independent of tumor region. These genes likely reflect core biological processes associated with tumor development that are common to both T1 and T2. In contrast, 697 genes were exclusively DE in T1 vs benign tissue, and 711 genes were DE only in T2 vs benign tissue, indicating distinct molecular features that differentiate tumor T1 and T2 from benign tissue and from each other **(Fig. 3B, Supp. Fig. 3C**).

Volcano plots **(Fig. 3C)** highlighted distinct expression patterns between benign, T1, and T2 tumor regions. To further explore the expression trends, we visualized the top 10 up– and downregulated genes in a heatmap across all the samples. Samples clustered distinctly by conditions, clearly showing higher similarity between T1 and T2 compared to benign (**Fig. 3D, Supp. Fig 3C**). Both T1 and T2 showed dysregulation of several key genes compared to the adjacent benign tissue, including COL2A1 (Collagen Type II Alpha 1 Chain) and cell cycle/proliferation markers such as BUB1B (mitotic checkpoint serine/threonine kinase B), CDCA8 (Cell division cycle associated 8), and PKMYT1 (Protein kinase, membrane associated tyrosine/threonine 1). Interestingly, some developmental transcription factors and regulators, such as SIX2 and EYA1 (EYA transcriptional coactivator and phosphatase 1) (Pierce et al., 2014, Petrosyan et al., 2023, Drake et al., 2009), were notably upregulated in T2 compared to benign tissue (see also IHC staining for SIX2 in **Supp. Fig. 1**), whereas T1 showed only modest and more variable expression levels. A marker of embryonic nephron progenitor identity of Wilms tumor, SALL1 (spalt like transcription factor 1), was found uniquely upregulated in T2-vs-benign and T2-vs-T1. Further we validated these findings using TARGET pediatric Wilms tumor dataset (samples n=130), which indicated that our tumor samples share a similar gene expression pattern with the TARGET dataset, for both the upregulated and downregulated genes. Pearson correlation between the TARGET average expression profile and the average expression of DE genes was 0.63 for T1 upper tumor and 0.67 for T2 lower tumor (**Supp. Fig. 4**) (Cerami et al., 2012).

The pathway enrichment analysis of differentially expressed genes of both tumor regions compared to benign showed significant (adjusted p-value (Benjamini–Hochberg FDR) < 0.05) activation of cell cycle related pathways, including G2/M checkpoint, E2F targets and mitotic spindle, suggesting cell cycle abnormalities and increased proliferation, supported by our high Ki-67 staining (see Supp. Fig. 5A). T1 exhibited a broader range of enriched pathways, indicating a more extensive dysregulation compared to T2 (**Fig. 3E**).

### Patient-derived cancer cell cultures recapitulate the parent tumor tissue regions

We established PDCs from both T1 and T2 to study the functional heterogeneity of these geno– and phenotypically divergent tumor regions, and the consequences for drug responses. Different cell culture conditions were tested: 2D PDCs were cultured either on Matrigel-coated (Mgel) plates or on irradiated mouse fibroblast (IR) feeder layers (Saeed et al., 2018, Liu et al., 2012), or in 3D in Mgel. All PDC models were validated for their representativity of original tumor tissues by both whole exome sequencing and immunostaining. Sequencing revealed a comparable mutational landscape between the tumor tissues and their corresponding PDCs, including shared mutations in *ASXL1* and *F7*, *TP53* mutation restricted to T1, as well as *ARID1A* and *KMT2D* variants specific to T2 (see **Fig. 2A**). CNAs, including gains on chromosomes 2, 8p, 13, 19q, and 20q, were identified in both T1 tissue and its derived PDCs, supporting the use of PDCs as real-time models for studying drug vulnerabilities associated with these alterations (see **Fig. 2E-F)**. In contrast to T1, both T2 tissue and corresponding PDCs displayed a largely copy-neutral profile.

Phenotypic characterization of PDCs (2D, Matrigel-grown, passage 3) was performed using a miniaturized multiplexed immunostaining platform in 384-well plates followed by high-content confocal imaging (**Supp. Fig. 5**). Marker expression in PDCs was compared with the corresponding immunohistochemical staining patterns observed in the original tumor tissues. A broad panel of epithelial, mesenchymal, proliferation, and signaling markers demonstrated retention of key phenotypic features in both T1 and T2 PDCs, consistent with the corresponding tumor tissues **(Supp. Fig. 5A**).

Consistent with the tissue analyses, T1 PDCs exhibited increased expression of WT1, a hallmark marker of Wilms tumor, whereas WT1 expression was markedly lower in T2 PDCs. WT1 expression was further validated by immunoblotting, which demonstrated substantially higher WT1 protein levels in T1 compared with T2 PDCs (**Supp. Fig. 5B**).

The IHC-staining pattern for nuclear localization of TP53 in T1, and low/absent nuclear TP53 in T2 (see **Fig. 2B**) was recapitulated in the matching PDCs, detected by immunofluorescence staining. (**Fig. 2C**). This is consistent with nuclear accumulation of TP53 in T1 whereas wild-type TP53 in T2 remains at low levels under basal conditions. Immunoblotting confirmed higher TP53 protein levels in T1 compared with T2 PDCs (**Fig. 2D**).

Overall, the PDCs retained key phenotypic and genetic characteristics of the T1 and T2 tumor tissues from which they were derived, as demonstrated by exome sequencing, immunostaining, and immunoblotting.

### *Ex vivo* drug sensitivity testing of PDCs reflects the underlying genomic and transcriptomic heterogeneity

The PDCs were subjected to *ex vivo* DSRT of hundreds of approved and investigational compounds (**Fig. 4A-B; Supp. Fig. 6; Supp. Table 2**) as well as to combinational therapies selected by clinicians treating the patient (**Fig. 4C, Supp. Table 3**). DSRT assays were performed using early-passage 2D PDCs cultured either on Mgel wells or on irradiated IR feeder layers, as indicated. As organotypic 3D PDC cultures did not achieve sufficient growth after the first passage (p1), the measurement of their drug sensitivity was not feasible. DSRT was performed first for PDCs at p1 with a miniaturized drug library of 116 compounds (FS2A; **see Supp. Table 2 for compound list and Supp. Table 4 for DSRT results**) and continued in subsequent passages with a comprehensive library of 528 compounds (FO5A, IR; T1 p6 and T2 p3, **Supp. Table 5**). The drug-tested PDCs correspond to the same early-passage cultures that were sequenced by WES **(Fig. 2, Supp. Fig. 2)**, ensuring molecular comparability across assays as well as the genotypic representativity of PDCs of tumor tissues.

**Figure 4.**
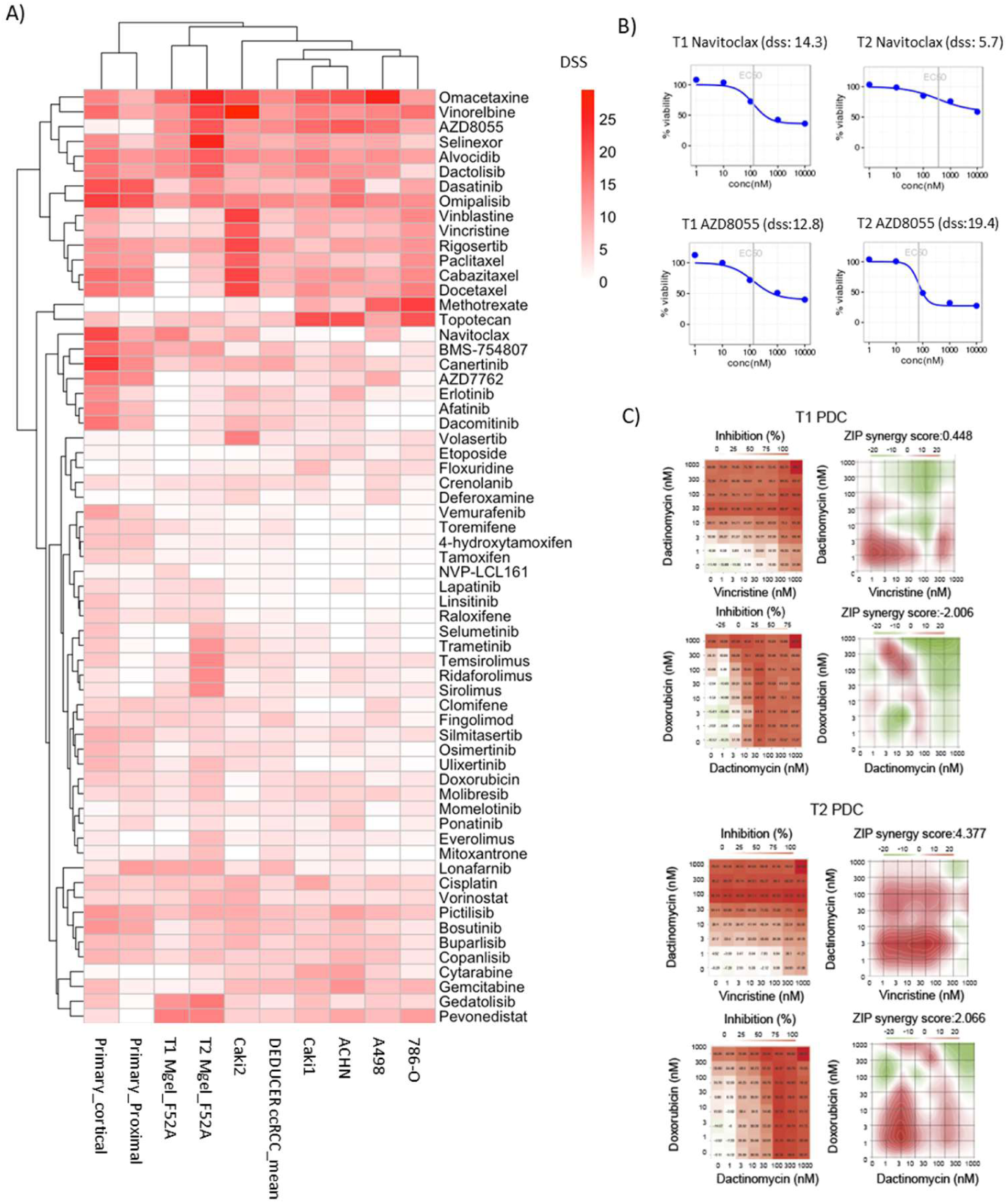
Drug sensitivity and resistance in T1 and T2 PDCs. **A)** Heatmap of DSS values across a panel of chemotherapeutic agents and targeted inhibitors in T1 and T2 PDCs along with mean DSS from DEDUCER ccRCC samples (n=18, Feodoroff et.al., 2026). Agents tested across all samples with the DSS > 5) were included in the heatmap. See also Supplementary Tables 4 and 5 for all T1 and T2 results **B)** Representative dose–response curves illustrating T1-selective sensitivity to the BCL-2 inhibitor navitoclax and T2-selective sensitivity to the PI3K/mTOR inhibitor AZD8055 **C)** Drug synergy analysis in T1 and T2 PDCs. Heatmaps show percent inhibition (left) and ZIP synergy scores (right) for vincristine-dactinomycin and doxorubicin-dactinomycin combinations. For inhibition, a sequential color scale was used, whereas ZIP synergy was visualized using a diverging color scale centered at zero, with positive values indicating synergy and negative values indicating antagonism.

We compared the drug sensitivity profiles of the T1 and T2 PDCs with those of healthy kidney cells (primary cortical and proximal), ccRCC cell lines (Caki-1, Caki-2, ACHN, A-498, and 786-O; DSRT with FO4B library), and with DEDUCER ccRCC samples (n = 18, Feodoroff et al., 2026). This analysis revealed that T1 and T2 exhibited distinct drug sensitivity profiles compared with all reference groups (**Fig. 4A)**. Overall, it was observed that the T2 PDCs were more sensitive to treatments compared to the T1 PDCs; for example, on average 40 compounds exceeded drug sensitivity score (DSS) ≥ 10 in T2 versus 31 in T1. Despite this overall pattern, distinct tumor_region-specific drug sensitivities were observed for both samples across several clinically approved agents. The TP53-mutant T1 PDCs showed increased sensitivity to selected compounds, particularly BCL-2 pathway inhibitors, including navitoclax (mean DSS T1 = 11.2, T2 = 2.1, **Fig. 4B**) and AT-101 (mean DSS T1 = 18.4, T2 = 14.2), as well as to the topoisomerase I inhibitor topotecan (mean DSS T1 = 10.5, T2 = 6.8) and the STAT3 pathway inhibitor napabucasin (mean DSS T1 = 18.7, T2 = 12.9) (**Fig. 4A6; Supp. Tables 4 and 5**). **; Supp. Fig.**

In contrast, T2 cells displayed consistently higher sensitivity to e.g. PI3K/mTOR pathway inhibition, as exemplified by dactolisib (dual PI3K/mTOR inhibitor; mean DSS T1 = 12.3 vs T2 = 17.6), AZD8055 (mTOR inhibitor; mean DSS T1 = 13.6 vs T2 = 18.8, **Fig. 4B)**, and omipalisib (dual PI3K/mTOR inhibitor; mean DSS T1 = 13.3 vs T2 = 17.3). A similar T2-biased sensitivity pattern was observed for the multi-targeted tyrosine kinase inhibitor dasatinib (mean DSS T1 = 5.1, T2 = 14.5) and taxanes (e.g., docetaxel T1 = 3.8 vs T2 = 13.6), indicating divergent pathway dependencies between the tumor regions.

Despite the *MET* copy-number gain in T1, functional testing did not support a strong MET dependency. Among MET inhibitors in the DSRT panel, capmatinib, crizotinib, and cabozantinib were tested, but all PDC models remained insensitive (DSS < 1.2).

For the combination testing in PDCs, the oncologists selected 12 pairs of clinically relevant drugs, typically used in metastatic Wilms tumor or in clinical trials for relapsed solid tumors (e.g., sarcomas) after first-line therapy (**Fig. 4C, Supp. Table 3**). Synergy between drug pairs was quantified using the zero-interaction potency (ZIP) model, where positive ZIP values indicate synergistic interactions (Yadav et al., 2015). In T1 PDCs, modest synergy was observed only for a limited number of combinations, with the strongest effect detected for doxorubicin + cyclophosphamide (ZIP = 1.77). In contrast, T2 PDCs showed broader and markedly higher synergistic responses. The combination of vincristine + dactinomycin produced the highest synergy (ZIP = 4.38) (**Fig. 4C**). A combination of dactinomycin + doxorubicin also showed measurable synergy in T2 (ZIP = 2.33). Accordingly, the patient received post-operative chemotherapy with vincristine, doxorubicin, and dactinomycin, showing ongoing relapse-free survival for more than 5 years. These drugs were also tested individually on the PDCs. Single-agent testing revealed partial to strong efficacy for all three vincristine (mean DSS T1 = 7.6, T2 = 9.3), doxorubicin (mean DSS T1 = 5.4, T2 = 7.9), and dactinomycin (mean DSS T1 = 29.9, T2 = 24.7).

To evaluate the functional relevance of the clinically actionable alterations identified through the iMTB analysis, we next compared the iMTB/PCGR-predicted biomarker–drug associations with the *ex vivo* responses obtained from the DSRT analysis. 20 of the 35 predictive biomarker drugs/drug sets were covered by our DSRT analysis (**Supp. Table 6**). Of note, while some of the drugs predicted to be effective based on T1 genetic biomarkers did achieve reasonable responses (e.g. the six tested anthracyclines ∼ TOP2A gain had a mean DSS of 7.9), average *ex vivo* response to these drugs was low (mean DSS across all of them: 4.1) and there was further no significantly stronger response for i) sensitivity vs resistance-predicted drugs within T1 (Wilcoxon p=1, though only Tamoxifen was resistance-predicted) (**Supp. Fig. 7A**) or ii) sensitivity-predicted drugs in T1, which did harbor the respective predictive biomarkers, vs the same drugs in T2, which did not harbor these biomarkers (Wilcoxon p=1, **Supp. Fig. 7B**). The most effective drugs identified from the DSRT were not associated with any genomic biomarker in either T1 or T2, confirming the added value of the *ex vivo* functional testing.

### HALLMARK gene-set analysis reveals that copy-number gains and differential expression are predictive of drug response

Because single molecular features often have limited predictive power for drug response (Madani Tonekaboni et al., 2020), we next integrated DSS values of drug testing, somatic mutations, CNAs, and differential gene expression quantified as RNA log₂ fold change (log₂(FC) of tumor vs benign) across MSigDB HALLMARK gene sets (**Fig. 5 A**). This multi-layer analysis revealed a substantial degree of intra-tumor heterogeneity.

**Figure 5.**
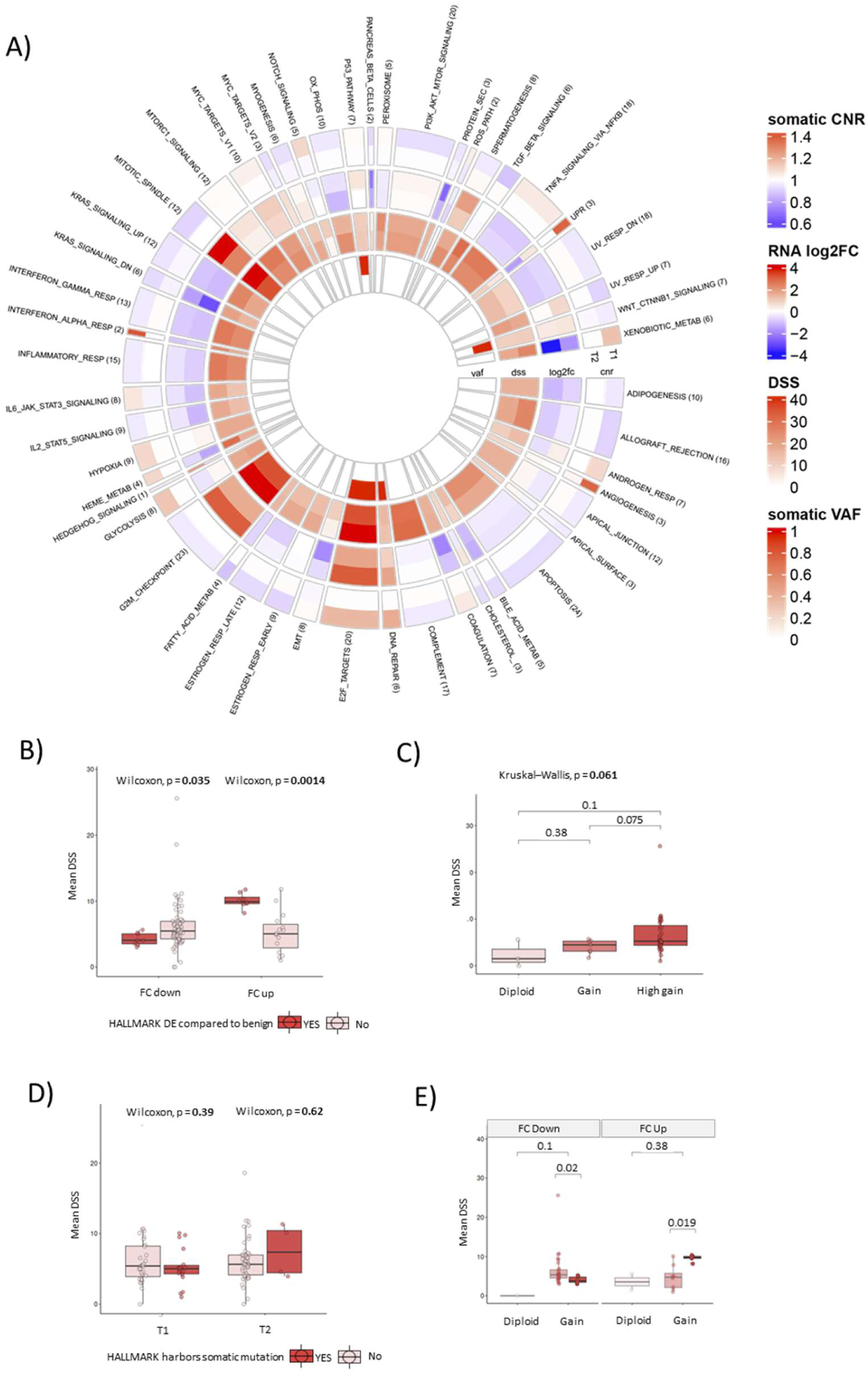
**A**) Multi-layered circular heatmap visualizing multimodal tumor profiling across MSigDB HALLMARK gene sets. Each ring represents different type of molecular data across two tumors (T1 and T2): (from outer to inner) Somatic copy number ratio (CNR), log 2-fold change in RNA expression (RNA log₂(FC), tumor vs benign), drug sensitivity score (DSS) and somatic variant allele frequency (VAF). The circle plot shows summary values for those HALLMARK gene sets targeted by at least one drug in the DSRT screens: median CNR and log₂(FC) and maximum VAF and DSS per gene set (see also **supp. Fig. 7D-E** for gene level data). The number in parentheses after the HALLMARK name indicates the number of distinct genes in each pathway which were targeted by the DSRT. **B)** Comparison of mean DSS values for HALLMARK gene sets stratified by differential expression relative to benign tissue. Gene sets with downregulated expression (FC down) exhibit significantly lower DSS (Wilcoxon p = 0.035), whereas upregulated gene sets (FC up) display significantly higher DSS (Wilcoxon p = 0.0014). **C)** Association between drug sensitivity (mean DSS) and maximal copy-number (CN) status within each HALLMARK gene set in the T1 tumor region. Diploid indicates gene sets without any CNAs; gain indicates at least one low-level CN gain (but no high-level amplifications); and high gain indicates gene sets with at least one high-level CN amplification. DSS increased stepwise from diploid → gain → high gain (Kruskal–Wallis p = 0.061). **D)** Mean DSS values for HALLMARK gene sets grouped by the presence or absence of somatic mutations in T1 and T2. No significant differences were detected in either region (T1: Wilcoxon p = 0.39; T2: Wilcoxon p = 0.62), indicating that mutation status alone does not predict ex vivo drug sensitivity in these gene programs. **E)** Joint representation of maximal CN status and differential expression (FC down / FC up) within HALLMARK gene sets in T1. Solid colors indicate gene sets with significant differential expression (DE); transparent colors represent non-DE gene sets. The highest average DSS was achieved for HALLMARK gene sets with co-occurring high-gain CNA and increased gene-expressions.

Consistent with the transcriptomic HALLMARK enrichment results (**Fig. 3E)**, the G2M checkpoint, E2F targets, and mitotic spindle gene sets emerged as the strongest and most consistently dysregulated HALLMARKs in both tumor regions (**Fig. 5A**). In these HALLMARKS, higher expression levels and variation in genomic alterations tended to co-occur with increased DSS, particularly in T1, suggesting region-specific dependencies on proliferative and cell-cycle–associated gene programs. Some HALLMARKs displayed tumor-compartment–specific signatures. For example, the NOTCH signaling gene set showed copy-number gains, transcriptional upregulation, and elevated DSS values in T1, whereas T2 exhibited minimal transcriptional activation and correspondingly low drug sensitivity for agents targeting this pathway, indicating functional NOTCH dependence unique to T1.

Across all HALLMARKs targeted by the DSRT, both differential expression and copy number status were predictive of *ex vivo* drug response, whereas somatic mutation status was not (**Fig. 5B-E**). Specifically, higher DSS values were achieved for i) HALLMARKS harboring one or more CNA gains (Kruskal–Wallis p=0.061, **Fig. 5C**), with higher level amplifications associated with even higher DSS (Wilcoxon p=0.075) and ii) HALLMARKS with significantly higher expression in the tumor compared to benign (Wilcoxon p=0.0014, **Fig. 5B**). Lower DSS values were achieved by HALLMARKS with significantly lower expression in the tumor compared to benign (Wilcoxon p=0.035, **Fig. 5B**), possibly due to a predominance of inhibitors in the DSRT.

Within T1 we also observed higher DSS for HALLMARKS with CNA gains and co-aligned significantly higher gene expression (Wilcoxon p=0.019, **Fig. 5E)**, whereas DSS was reduced for HALLMARKS with CNA gains but co-occurring significantly reduced expression (Wilcoxon p=0.020). While T2 was excluded from this overlap analysis as it did not harbor any CNAs in any of the investigated HALLMARKs, T1 harbored at least one CNA in each of the differentially expressed ones (i.e. no differential expression observed in the absence of CNAs), suggesting again different mechanisms underlying the molecular deregulation of these two samples.

Although these same patterns of differential expression, copy number, mutation status and DSS were observed also at the level of individual targeted genes, the pattern was stronger and statistical significance greater at the HALLMARKs level (see gene-level plots in **Supp. Fig. 7 D-E**).

## DISCUSSION

In this study, we investigated an adult Wilms tumor using a pathway-centric functional precision medicine approach that integrates histopathology, multi-region multi-omics, and *ex vivo* drug testing of PDCs, together with benchmarking our transcriptomic findings against the TARGET Wilms Tumor dataset. To our knowledge, this represents the first systematic *ex vivo* drug profiling of PDCs derived from an adult Wilms tumor, together with other omics data, revealing both investigational and approved therapeutic opportunities not apparent from genomics alone. We show that, although intratumoral heterogeneity spans pathological, genomic, transcriptomic, and functional layers, shared and targetable pathway programs are present —most notably the G2/M checkpoint, E2F targets, and mitotic spindle. These data highlight the value of combining multi-region molecular profiling with functional assays to identify actionable dependencies in this malignancy and to refine therapy selection beyond single-gene biomarkers.

We observed marked histological heterogeneity between spatially distinct regions of the studied Wilms tumor. Both regions fulfilled diagnostic criteria for Wilms tumor yet differed in cellular composition (relative proportions of blastemal, epithelial, and stromal elements), proliferative indices, and immunophenotypic marker expression. These observations align with the established heterogeneity of Wilms tumor (Cresswell et al., 2016; Škarda et al., 2025), including local anaplasia and region-specific biomarker patterns. Such compartmentalization has direct implications for treatment response and biomarker interpretation, supporting multi-region sampling and integrated profiling to guide therapy in adult Wilms tumor.

We found several actionable, clinically relevant alterations in the tumor areas through WES, most notably the TP53 mutation in T1, alongside CNAs involving genes linked to cell-cycle regulation and oncogenic signalling. Spatially resolved genomics in Wilms tumor has earlier revealed substantial intra-tumor heterogeneity and branched evolution in pediatric cohorts, with region-specific drivers and local anaplasia linked to outcome (Cresswell et al., 2016). Consistent with this, this adult case of Wilms tumor harbored a truncal ASXL1 frameshift across both regions, alongside compartment-specific alterations—TP53 and copy-number gains in T1, and ARID1A/KMT2D truncations with a copy-neutral genome in T2—together with distinct ATM mutation patterns. While ASXL1 is not among the most frequent Wilms tumor genes, it has been recurrently implicated in pediatric series (Gadd et al., 2017) and may represent an early chromatin remodeling lesion that primes nephrogenic programs and global transcriptional dysregulation. Against this background, region-specific genetic alterations segregated along canonical Wilms tumor axes: DNA damage response disruption (TP53, ATM) and chromatin/enhancer remodeling (ARID1A, KMT2D, SWI/SNF and H3K4 methylation), both recognized in pediatric disease (Gadd et al., 2017) and here in an adult tumor with spatial resolution.

*Ex vivo* drug testing performed for pheno– and genotypically representative PDCs provided critical functional insights that could not be inferred from genomics alone. Combination drug testing further revealed region-specific synergistic responses, with broader and stronger synergy observed in T2.

In T1, co-alteration of ATM and TP53 suggests disruption of the same DNA damage response and tumor suppressor axis, potentially impairing checkpoint activation, p53-mediated apoptosis, and DNA repair. This may be explained by the observed sensitivity to agents that exploit replication stress and apoptotic priming, including topoisomerase I inhibitors and BCL-2 family inhibition. Moreover, ATM– and/or TP53-deficient tumors may become more dependent on compensatory DDR pathways, including ATR–CHK1–WEE1 signaling, supporting the rationale for testing DDR-targeted combinations such as ATR, CHK1, WEE1, or PARP inhibition (Jette et al., 2019; Drew et al., 2025).

In contrast, T2 exhibits an ATM “double-hit” without TP53 mutation, alongside ARID1A/KMT2D loss-of-function. These features suggest checkpoint fragility and enhancer reprogramming within a copy-neutral genome—hallmarks associated in other cancers with reliance on mitotic and replication stress pathways, such as those targeted by microtubule agents or ATR/CHK1 inhibitors. This “wiring” could potentially underlie the broader single-agent responses and stronger combination synergies we observed in T2. Importantly, these pathway-informed combinatorial vulnerabilities may inform hypothesis generation around therapeutic prioritization in adult Wilms tumor, where treatment-related toxicity remains a major clinical challenge. However, the genomic biomarkers alone were insufficient to predict functional drug responses, in line with growing evidence that single molecular alterations often have limited predictive power, particularly in heterogeneous tumors.

Adult Wilms tumor is frequently presumed to differ biologically from pediatric disease; however, direct molecular comparisons have been limited by the scarcity of adult cases. Transcriptomic profiling revealed that, despite regional genomic differences, both tumor regions showed activation of proliferative and cell-cycle–associated pathways, including G2/M checkpoint, E2F targets, and mitotic spindle programs. The observation that these cell-cycle–associated pathways are activated both in this adult case and in the TARGET pediatric cohort suggests that – despite genetic and clinical differences-that core proliferative programs may be conserved across age groups. This pathway-level activation was more informative than individual gene alterations for predicting drug responses, suggesting that shared transcriptional programs may represent robust therapeutic targets even in the presence of genetic heterogeneity. Benchmarking our findings against the TARGET Wilms Tumor cohort mitigates the limitations inherent to an n=1 study and supports the generalizability of the identified pathway-level dependencies.

Clinically, our data refine and extend existing Wilms tumor treatment strategies. Standard pediatric backbones (vincristine, dactinomycin, doxorubicin) showed *ex vivo* activity and synergy, co-aligning with the patient’s favourable outcome after the radical nephrectomy. Additional therapeutic hypotheses emerged from functional profiling — topoisomerase I inhibitors and BCL-2 family inhibitors in the TP53/ATM-altered tumor region, and microtubule/mitotic agents in combination regimens in the ARID1A/KMT2D/ATM-altered, copy-number-neutral tumor region. While these region-specific drug sensitivities and combination synergies should be considered exploratory, they highlight biologically supportive vulnerabilities that justify further investigation. In contrast, the shared activation of proliferative and cell-cycle–associated pathways across spatially distinct tumor regions, supported by benchmarking against the TARGET cohort, represents a more robust finding of this study. These findings support multi-region sampling and integrated functional testing in adult Wilms tumor, where standardized adult guidelines are lacking and randomized evidence is unattainable due to extreme rarity.

Limitations of our study include the n = 1 design and *ex vivo* model constraints (absence of microenvironmental and pharmacokinetic effects). We partially mitigated these by benchmarking transcriptomic signatures against the TARGET cohort (n = 130) and demonstrating similar pathway activation pattern. In addition, we compared the drug testing results of the studied tumor PDCs to ccRCC PDC cohort (n = 18) as well as cancer cell lines (n = 5). Prospective application of this pathway-centric, multi-region functional approach across additional adult cases and high-risk pediatric tumors will be essential to validate generalizability, refine biomarker panels (e.g., combined expression/CNA scores for G2/M/E2F/mitotic spindle), and prioritize rational combinations for clinical testing in Wilms tumor.

Altogether, these findings support a pathway-centric, multi-region functional precision medicine strategy that characterises druggable biological heterogeneity and driver signaling pathways in rare cancers, accelerates the translation of robust predictive biomarkers into clinical practice, and informs individualized therapy options.

## MATERIALS AND METHODS

### Patient tissue sampling to PDCs and PDC cultures

The sectioned tissue was sent to routine tissue processing and paraffin embedding, followed by H&E staining for microscopic evaluation. Before formalin fixation, two pieces (app. 1 cm x 1 cm each) from macroscopically distinct areas of the tumor were visually sampled by a pathologist for subsequent PDC cultures. Hereafter, these pieces are referred to as tumor region 1 (T1; upper part of the kidney) and tumor region 2 (T2; lower part of the kidney). The tissue pieces were further divided for 1) tissue dissociation, 2) DNA isolation, and also 3) freshly frozen samples for further experiments. The pathologist confirmed the presence of tumor cells from tissue samples stained in the diagnostics lab with H&E.

Tumor tissues were dissociated with Tumor Dissociation Kit (Miltenyi, 130-095-929) with GentleMACS C tubes (Miltenyi) and GentleMACS Dissociator (Miltenyi). After dissociation, cells (app. 1-1.5 million cells/ 10 cm2 dish) were added to irradiated 3T3 feeder cells (Saeed et al. 2018) and on Matrigel (Corning 354234) coated dishes in F-medium. Complete F-medium contained (F-12:DMEM, v/v 3:1) DMEM (Gibco, 42430-025) and F-12 Nut mix (Ham) (Gibco, 21765-029) supplemented with Rock inhibitor (Y-27632) 10 µM (Enzo Life Sciences, ALX-270-333), Penicillin 10 U/ml: Streptomycin 10 µg/ml (Gibco, 15040-122), 5 % FBS (Gibco, 10270-106); 10 ng/ml EGF (Sigma, E9644), 0.4 µg/ml Hydrocortisone (Sigma, H088); 2,1 ng/ml Choleratoxin (Sigma, C8052), 24 µg/ml Adenine (Sigma, A2786) and 5 µg/ml Bovine insulin (Sigma, I0516). The cells were divided 1:2-1:3 once or twice per week and in addition to this, medium was changed once per week.

### Culture of primary renal epithelial cell lines and renal cancer cell lines

Commercially available primary renal cortical epithelial cells (Pan-Cytokeratin (+), TE-7 (-), normal, human; HRCE, ATCC, PSC-400-011) and primary renal proximal tubule epithelial cells (Pan-Cytokeratin (+), y-glutamyltransferase-1 (GGT-1) (+), TE-7 (-), normal, human; RPTEC, ATCC, PCS-400-010) were cultured as in manufacturer’s protocol in Renal Epithelial Cell Basal medium (ATCC, PCS-400-030) supplemented with Renal Epithelial cell growth kit (ATCC, PCS-400-040). Medium was changed every 2-3 days and cells divided 1:2 once a week.

Renal cancer cell lines, 786-O _[MPM1]_ (ATCC) and ACHN (NCI) were cultured in RPMI1640, Caki-2 (DSMZ) cells in McCoy’s 5A and A-498 (DSMZ) cells in MEM medium, and all were supplemented with 10 % FBS, Penicillin 10 U/ml: Streptomycin 10 µg/ml and L-glutamine. Caki-1 (DSMZ) cells were cultured in 20% FBS in McCoy’s 5A medium supplemented with Penicillin-Streptomycin and L-glutamine. Media were changed 2-3 times per week and confluent cultures split 1:3-1:6 once or twice per week. All cell lines were authenticated using Promega GenePrint24 system at Genomics Unit of Technology Centre (Institute for Molecular Medicine Finland-FIMM, HiLIFE, University of Helsinki, Finland).

### Immunophenotyping of tissues and PDCs

Clinical immunohistochemical (IHC) staining of the paraffin-embedded tissue samples was performed at HUSLAB using diagnostically validated antibodies targeting various renal cancer and Wilms tumor markers. The stainings were evaluated by an experienced pathologist (TM), and slides were imaged in the research laboratory using Pannoramic P250 Flash II whole-slide scanner (3DHistech, Hungary).

FFPE blocks of diagnostic material—including benign tissue, T1 tumor tissue, and T2 tumor tissue—were obtained from the Helsinki Biobank. The performed additional IHC stainings, including antibody suppliers, catalogue numbers, and dilutions, are listed in **Supp. Table 7**.

For the chromogenic IHC, slides were stained as in Blom et al. (2017). Briefly, the in-house IHC protocol was used for selected antibodies (**Supp. Table 7**), while some of the stains were combined using multiplexed immunostaining (mIHC) protocol (Blom et al., 2019) to further study the possible co-localization in the tissue samples. The immunohistochemical panel 1 included antibodies against carbonic anhydrase IX (CAIX), pan-epithelial markers (pan-cocktail), alpha-methyl coenzyme A racemase (AMACR), cytokeratin 7 (CK7), Ki-67 and vimentin. The multiplex panel 2 included CD3, CD163, pan-keratin, DAPI, CAIX and pan-cocktail. The mIHC stainings were imaged with Axio Scan.Z1 (Zeiss, Germany) with a 20x objective (NA 0.8) at the Digital Pathology (DP) core unit at FIMM.

The PDCs were seeded to 384-well plates (Corning, #3764), grown for 2 days to be subconfluent, fixed with 4% paraformaldehyde, incubated for 20 minutes at room temperature (RT) and washed with PBS by plate washer (Biotek, EL406). Cells were permeabilized with 0.3% Triton X-100 in PBS for 10 minutes and blocked with 3% BSA in PBS for an hour. Primary antibodies (described in the **Supp. Table 7)** were diluted in 1% BSA/0.1% Tween-20 in PBS and added to the cells, followed by incubation for 1 h at 37 °C. After washing with PBS, secondary antibodies Alexa Fluor 488 anti-rabbit (Invitrogen, A11008) and Alexa Fluor 568 anti-mouse (Invitrogen, A11004) were applied at a final dilution of 1:2000 together with Phalloidin 1:50 in 1% BSA/0.1% Tween-20 in PBS and incubated for 1 h at RT. Hoechst (10 mg/mL stock solution) was diluted 1:10,000 in 1% BSA in PBS and incubated with the cells for 5 minutes. All washing steps were performed using a plate washer (Biotek EL406).

Stained cells were imaged with Opera Phenix (Revvity) by using 20x water immersion objective (NA 1.0) (9 frames/well).

### Immunoblotting

Whole cell extracts were collected at the point of drug screening (FS2A), in RIPA buffer (Pierce, 89900) supplemented with proteinase inhibitor (Pierce, 88666) and phosphatase inhibitor (Pierce, 88667). The concentration of the isolated proteins was determined using BCA Protein Assay Reagent kit (Pierce, 23227). Tumor samples T1 and T2 were analyzed together with 786-O and OVCAR-3 (NCI-DTP) cell line controls and a ccRCC PDC control (Feodoroff et al., 2026). For each sample, 25 µg of protein was separated by SDS-PAGE on 7.5% gels (Bio-Rad, 4561025) and transferred to a polyvinylidene difluoride membrane (Millipore, IPFL00010) using the Mini Trans-Blot® Electrophoretic Transfer Cell according to the manufacturer’s protocol (Bio-Rad). To prevent nonspecific binding of primary antibodies, the membranes were blocked with Odyssey Intercept PBS blocking buffer (LI-COR Biosciences, 927-70001) for 60 min at room temperature and subsequently incubated with the primary antibodies at 4°C overnight. The primary antibodies, diluted 1:1000, were WT1 (Cell Signaling Technology, 83535), TP53 (Cell Signaling Technology, 9282) and E cadherin (BD Biosciences, 610182). Antibodies against GAPDH (Sigma Aldrich, G8795) and Beta-tubulin (Abcam, ab6046) were used as loading controls to confirm equal loading of the samples, and diluted 1:1000 and 1:500, respectively. After primary antibody incubations, membranes were washed four times for 5 min with 0.1% Tween-20 in PBS and incubated with fluorescently labeled secondary antibodies (LI-COR Biosciences, 926-32211, 926-68070) for 1h at room temperature. Finally, the membranes were washed four times for 5 min with 0.1% Tween-20 in PBS and once with PBS only and scanned with Odyssey LI-COR imaging system according to the manufacturer’s protocol.

### Drug sensitivity and resistance testing (DSRT)

DSRT assay was performed on PDCs (from passage 1 to passage 6), starting with a miniaturized drug library of 116 compounds (FS2A; see **Supp. Table 4** for compounds) and continuing with a whole library set of 528 compounds (FO5A; **see Supp. Table 5**), all in 5 concentrations on 384-well plates, prepared at the High Throughput Biomedicine Unit, FIMM, HiLIFE, UH, Finland. PDCs, originally established with IR feeder cells for the first passage, or on Mgel-coated dishes were detached and seeded to pre-drugged 384-well plates (Saeed et al. 2018) in 25 µl. The plates were incubated at 37°C with 5% CO^2^ for 72 hours prior to addition of CellTiter Glo (CTG 2.0, Promega, G9243) to wells in equal volume of 25 µl and detection of luminescence signals with microplate reader (PHERAstar FS, BMG Labtech). Combination screens of selected drugs were performed in a similar way (**Supp. Table 3**). The data was run through the quality analysis pipeline to confirm the robust Z prime (>0.70) for the assay (in-house analytics tool Breeze, Potdar et al., 2020). The drug sensitivity and resistance scores (DSS) were calculated for each compound to quantify drug efficacy across the tested concentration range (Yadav et al., 2014). Detailed target annotations and approval status of the screened compounds are provided in **Supp. Table 2.** Approval status was determined by querying openFDA, including FDA drug-label and Drugs@FDA data, the European Medicines Agency (EMA) medicines reports, ClinicalTrials.gov v2 API for the highest reported clinical phase, and the Open Targets Platform for maximum clinical phase and approval status. Mechanism-of-action annotations were retrieved from the Open Targets Platform and supplemented with ChEMBL mechanism-of-action records.

### DNA isolation and exome sequencing

Both tumor tissue DNA and blood germline DNA were isolated from fresh material upon sample arrival. The DNA from PDCs was extracted simultaneously to the first drug testing (passage 1 and 3). DNA extraction was performed from all samples using DNeasy Blood and Tissue (Qiagen, 69504) kit.

The exome sequencing was performed at FIMM Sequencing Unit. 50 ng of gDNA was processed according to the Twist Human Core Exome EF Multiplex Complete kit manual (Twist Bioscience, San Francisco, CA, USA). Adapters used for ligation were unique dual index (UDI) oligos with unique molecular barcodes (UMI) by IDT (Integrated DNA Technologies, Coralville, IA, USA). Library quantification and quality check was performed using LabChip GX Touch HT High Sensitivity assay (PerkinElmer, USA). Libraries were pooled to 8-plex reactions according to concentration. The exome enrichment was performed according to the Twist Human Core Exome EF Multiplex Complete kit (Twist Bioscience, San Francisco, CA, USA) manual, using Twist Core Exome probes and using Twist Human RefSeq Panel as spike in. Libraries were quantified for sequencing using the 2100 Bioanalyzer High sensitivity kit. Sequencing was performed with Illumina NovaSeq system using S4 flow cell with lane divider (Illumina, San Diego, CA, USA). Read length for the paired-end run was 2×151 bp.

Germline and somatic mutation analyses from whole-exome sequencing data was performed using the Illumina DRAGEN Bio-IT Platform v4.0.3 (https://support-docs.illumina.com/SW/DRAGEN_v40/Content/SW/FrontPages/DRAGEN.htm). Read alignment was carried out using the GRCh38 reference genome. Prior to alignment, reads were trimmed to remove low-quality bases (minimum quality score of 3), poly-G tails, and TruSeq adapter sequences.

Matched blood samples served as the normal reference to distinguish somatic alterations. Somatic variants were identified using the **‘**somatic mode**’** of DRAGEN’s small variant caller, which operates in tumor-normal mode, jointly analyzing tumor and normal samples. This method assumes that germline variants and technical artifacts are shared across both samples, while true somatic mutations are unique to the tumor sample. Further, ANNOVAR tool was used for the functional annotation of predicted somatic variants (Wang *et. al*., 2010). Gene copy number alterations (CNAs) were identified using the CopyCat tool (https://github.com/chrisamiller/copycat). For validation of our predictions we used data from Therapeutically Applicable Research to Generate Effective Treatments (TARGET) (https://www.cancer.gov/ccg/research/genome-sequencing/target) initiative, phs000471.

### Integrative Molecular Tumor Board (iMTB) reports

To identify the functionally and clinically relevant findings across data modalities, we used our in-house iMTB tool, developed on top of existing tools such as The Personal Cancer Genome Reporter (PCGR) for genomic biomarker annotation (Nakken et al., 2018).

### Transcriptomics

FFPE tissue samples were collected using three methods: (1) manual scraping of ROIs (81–144 mm²) from H&E-stained Superfrost Plus slides; (2) laser capture microdissection (LCM, Leica LMD 7) of unstained ROIs (∼11 mm²) from PEN membrane slides; and (3) LCM of deparaffinized slides stained with DAPI and concanavalin A for enhanced visualization (∼11 mm²). All samples were lysed in HTG EdgeSeq Lysis Buffer (HTG Molecular Diagnostics), with buffer volumes adjusted based on ROI area ((area in mm² / 11) × 50 μL). Lysis was performed at 56 °C for 20 min (850 rpm), followed by 95 °C for 15 min, then cooling on ice. After transferring 48 μL of lysate, DNase treatment (6 μL DNase + 6 μL buffer, 37 °C, 30 min) was applied, followed by addition of Biofluids Buffer (48 μL) and heating at 95 °C for 10 min. The samples were then frozen to –80 °, before pipetting into the HTG EdgeSeq barcoded sample plates.

RNA expression profiling was carried out using the HTG Transcriptome Panel on the HTG EdgeSeq platform, with sequencing on an Illumina NextSeq 500.

Differential gene expression analysis was performed using the **DESeq2** package (Love et. al., 2014). Tissue samples collected using three different methods (as described above) were used as biological replicates for T1, T2, and Benign sample groups. Genes were considered differentially expressed if they met the criteria of an adjusted *p*-value (Benjamini–Hochberg FDR) < 0.01 and a |log₂(FC)| > 2. To investigate pathway-level changes, we performed Gene Set Enrichment Analysis (GSEA) using the fgsea R package (https://github.com/alserglab/fgsea) with default parameters. Hallmark gene sets were retrieved from the MSigDB (Liberzon et.al., 2015) database via the msigdbr package. Enriched pathways were considered significant at an adjusted *p*-value (Benjamini–Hochberg FDR) < 0.05.

## AUTHOR CONTRIBUTIONS

Conceptualization, V.P., A.R., T.M.; Methodology, M.P., P.M., T.P., K.V., R-M.K., T.L., A.H., J.S., J.L., S.K., P.Ja., P.S., P.Jä., Investigation, M.P., P.M., R.K., M.M., T.L., J.S., K.M., T.P., J.L., R.K., V.P., T.M., E.R.; Writing – Original Draft M.P., R.K., T.L., V.P., P.M., Visualization, M.P., T.L:, R.K., V.P., P.M.; Funding Acquisition, O.K., V.P., T.M., A.R; Supervision V.P.

## ACKNOWLEDGEMENTS

This study has been supported by grants from Finnish Cultural Foundation, TEKES/Business Finland FiDiPro fellowship, Cancer Foundation, iCAN Digital Precision Cancer Medicine Flagship (iCAN), Finnish Cancer Center (FICAN), and Kymenlaakso Cultural Foundation. Helsinki Biobank and FIMM Digital Microscopy and Molecular Pathology Units are acknowledged. FIMM High Content Imaging and Analysis Unit (FIMM-HCA; EuroBioImaging), FIMM High Throughput Biomedicine Unit (FIMM-HTB), and FIMM Sequencing Unit at FIMM Technology Center (HiLIFE, University of Helsinki) and Biocenter Finland, are acknowledged for the technical expertise.

