## Supplementary Figures for "Pathway-centric multi-omics and functional precision medicine reveal shared drug vulnerabilities in heterogeneous adult Wilms tumor"

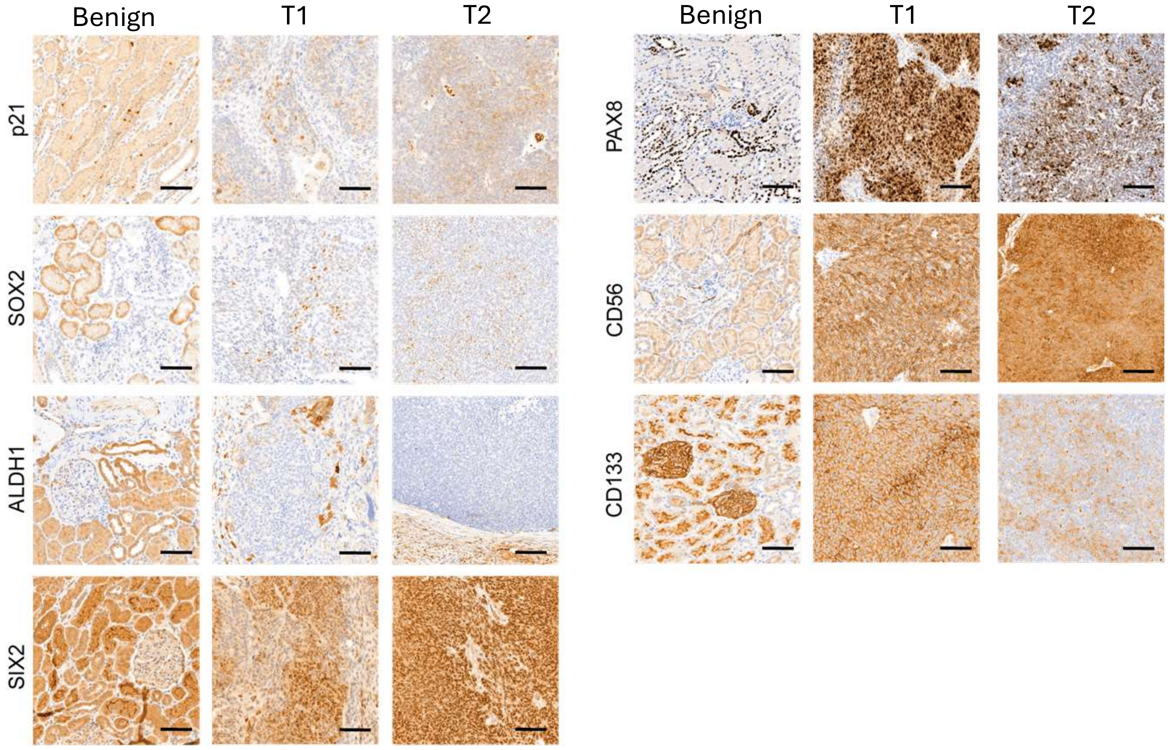


**Supplement Figure 1 (related to Figure 1).**
Immunohistochemical staining of benign, T1, and T2 tumor tissues for p21, SOX2, ALDH1, SIX2, PAX8, CD56 and CD133. Scale bars = 100 µm

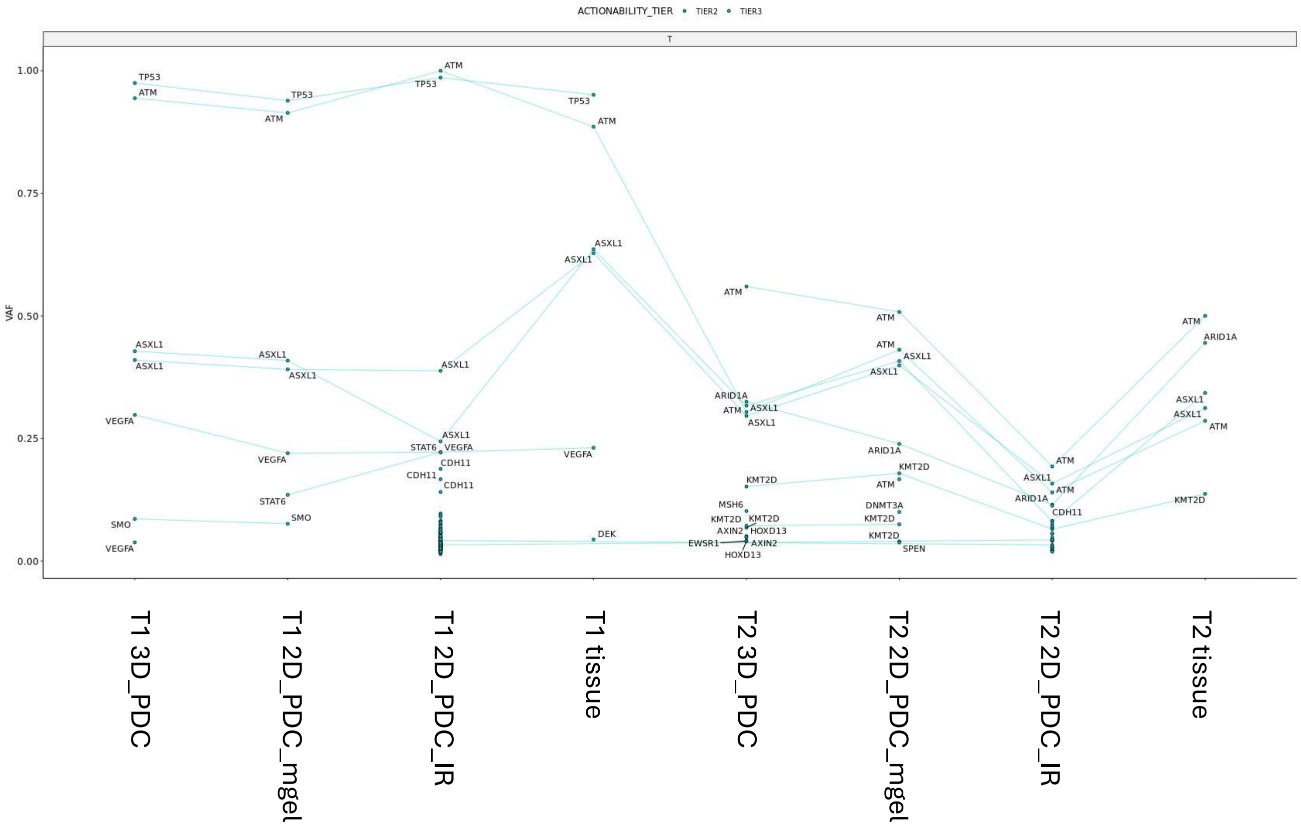


**Supplement figure 2 (Related to figure 2). Variant allele frequencies (VAFs) of clinically relevant somatic variants across Wilms tumor samples and derived PDCs.** VAFs of Tier 2–3 variants (no Tier 1 variants were observed). Each point represents an individual somatic variant, and lines connect identical variants across samples. Multiple unconnected points with the same gene symbol indicate distinct mutations within that gene.

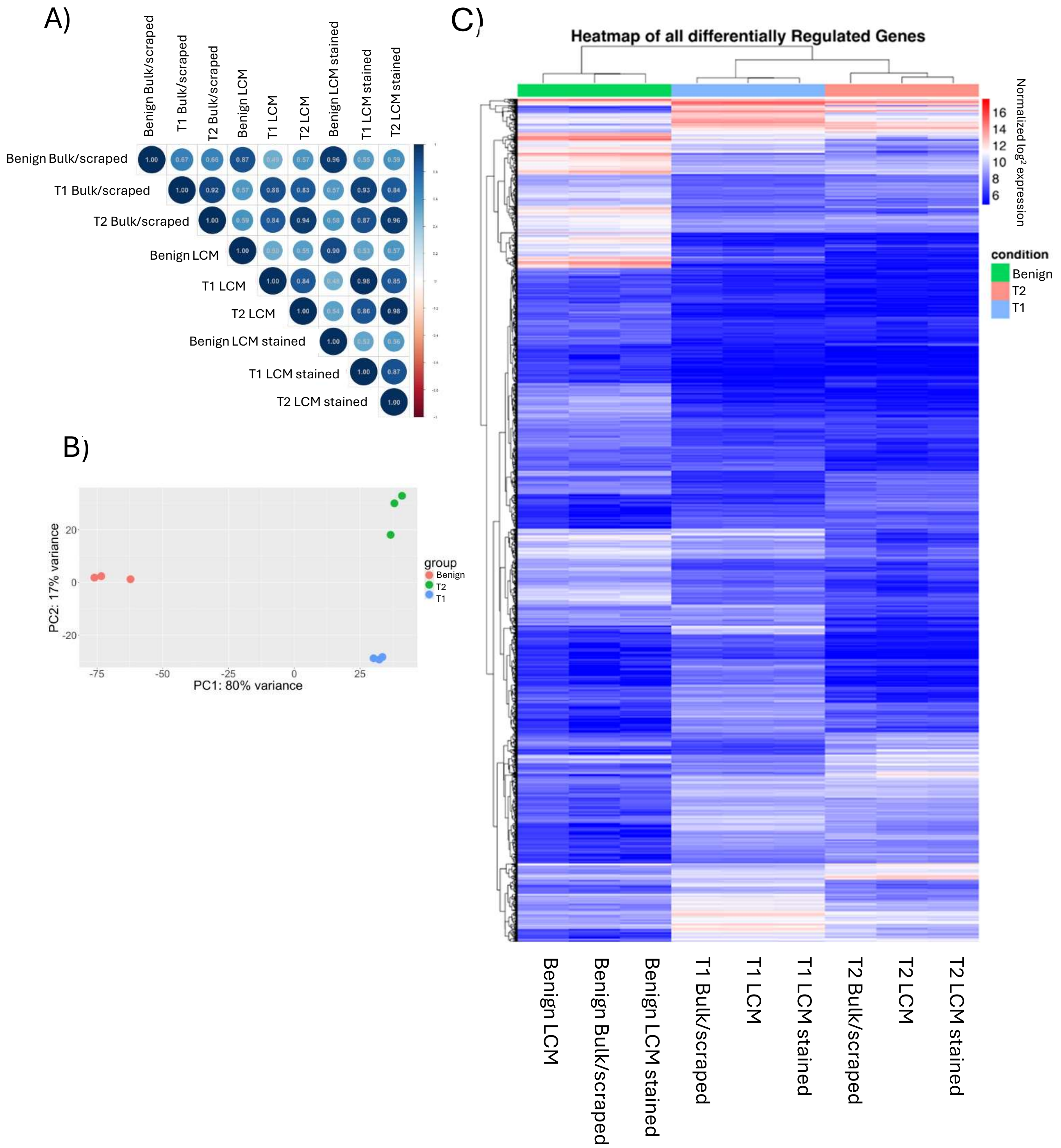


**Supplement Figure 3 (related to Figure 3). Transcriptomic profiling of benign tissue and two tumor regions T1 and T2. A)** Sample–sample correlation matrix showing pairwise Pearson correlation coefficients across all RNA-seq samples, including benign tissue, T1 and T2 tumor regions. Each condition (Benign, T1, T2) is represented by multiple technical preparations, including scraped/bulk-extracted RNA, laser-capture microdissection (LCM) from unstained sections, and LCM from stained sections. Darker blue indicates higher correlation, and red indicates lower correlation. **B)** Principal component analysis (PCA) of transcriptome-wide gene expression. PC1 (80% variance) separates T1 samples from benign and T2 samples, while PC2 (11% variance) captures variability primarily between benign and tumor groups. **C)** Global heatmap of all differentially expressed genes across benign, T1, and T2 samples. Rows correspond to genes and columns to individual samples from each extraction method (scraped, LCM, stained LCM). Hierarchical clustering demonstrates grouping by tumor region, illustrating consistent transcriptional patterns despite differences in sample preparation.

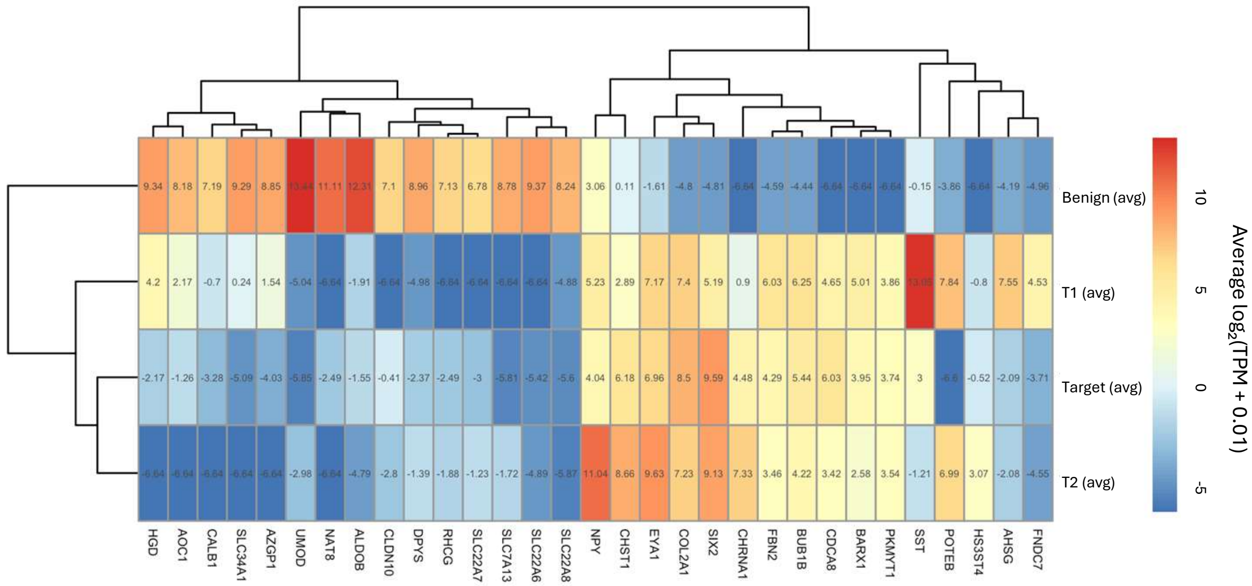


**Supplement Figure 4 (related to Figure 3)**. Heatmap comparing gene expression profiles of benign tissue, tumor region T1 and T2 with the cohort-level mean expression values from the TARGET Wilms Tumor dataset. Each row represents one sample group; benign (avg), T1(avg), T2 (avg), TARGET (mean), and each column corresponds to an individual gene. Values reflect Average $\log_2$(TPM + 0.01) expression levels.

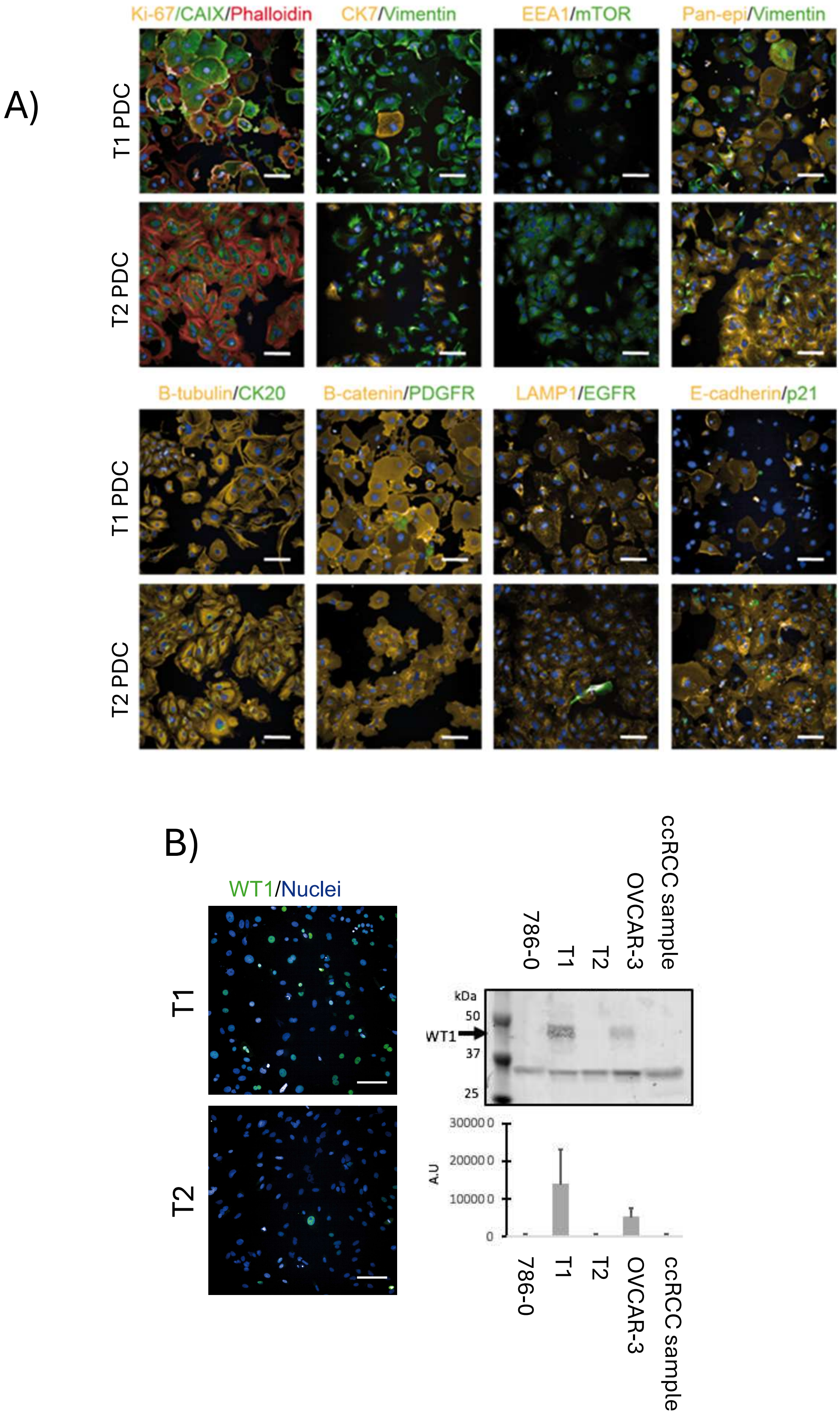


**Supplement Figure 5, related to figure 4. A)** Multiplex immunofluorescence (IF) staining of early-passage PDCs derived from regions T1 and T2. Representative confocal images showing marker expression patterns across multiple cellular compartments. Each marker combination is shown for both T1-PDC and T2-PDC. Nuclei are counterstained (blue). Scale bars = 100 µm. **B)** WT1 immunofluorescence staining and western blot analysis in patient-derived cells (PDCs) from T1 and T2. WT1 immunofluorescence staining and western blot analysis in T1- and T2-derived PDCs. WT1 protein levels differ between tumor regions despite the absence of known pathogenic WT1 mutations in either sample. Control cancer cell lines are shown for comparison. Scale bars = 100 µm.

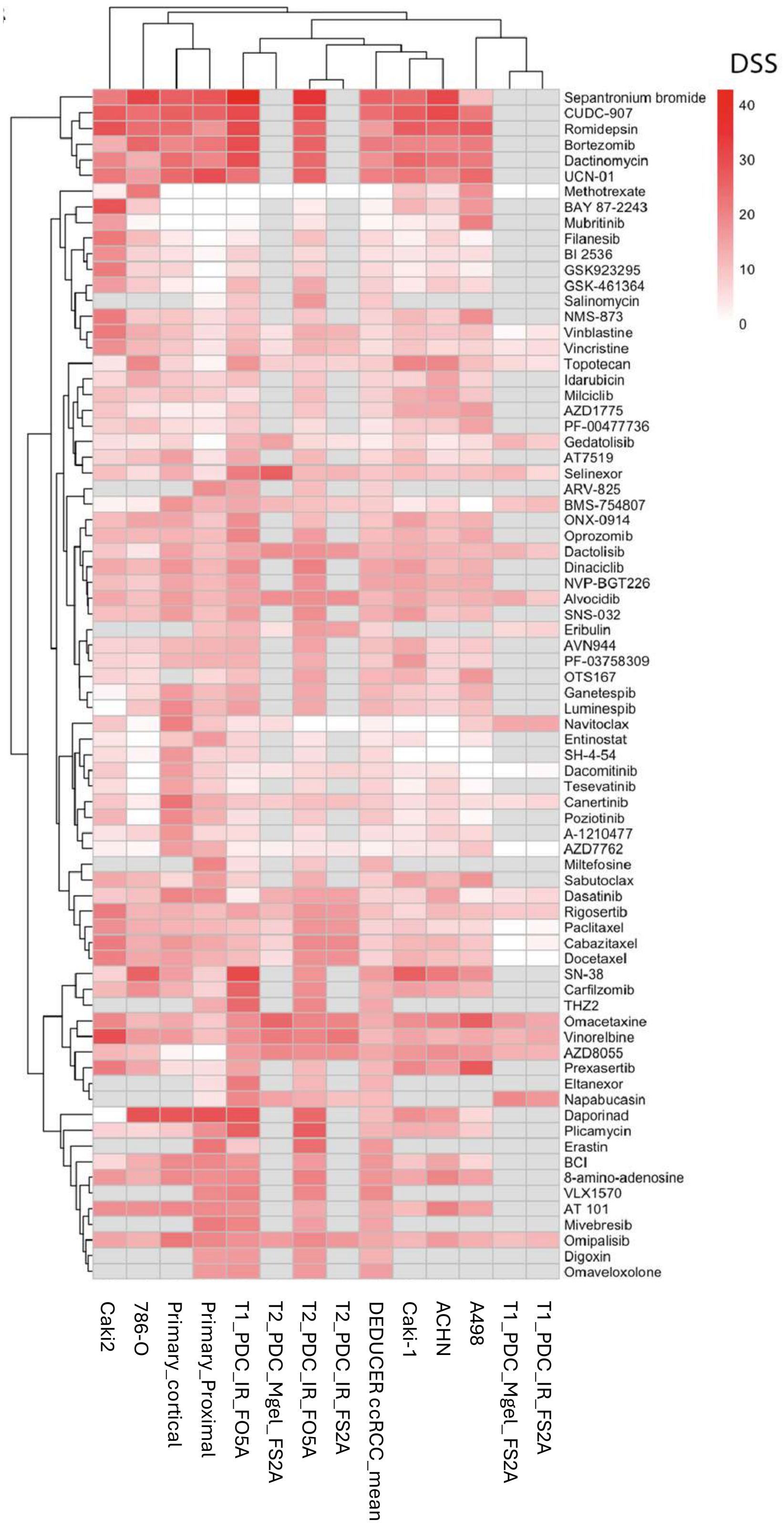


**Supplement Figure 6 (related to Figure 4).** Hierarchically clustered heatmap depicting DSS values for tested compounds in the FS2A/FO5A drug libraries. Rows represent individual compounds; columns represent replicate measurements or sample conditions. Color intensity corresponds to DSS magnitude (red = higher DSS; white = lower DSS). Grey color means not tested for that compound.

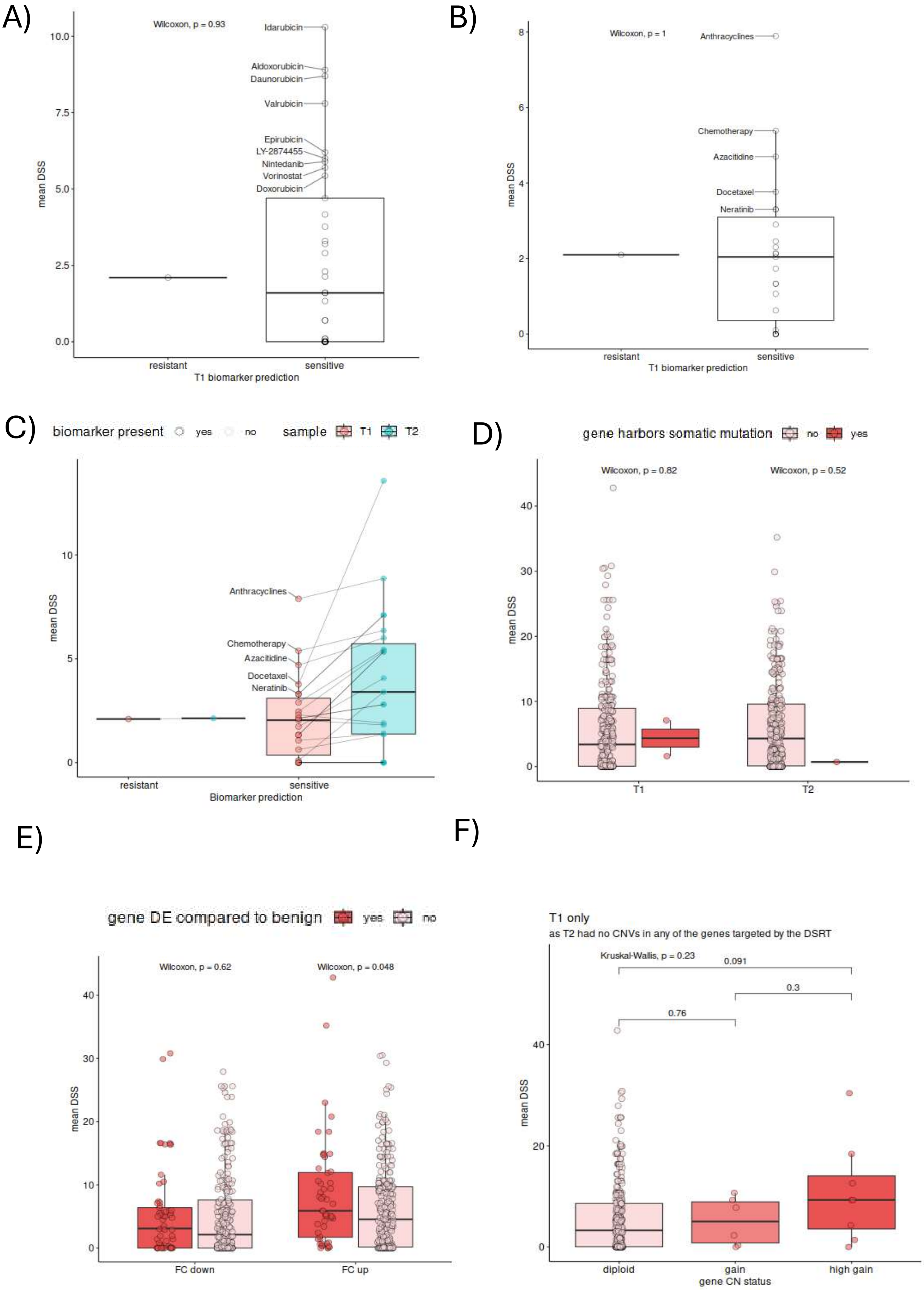
A)
Wilcoxon, p = 0.93
Idarubicin
Aldoxorubicin
Daunorubicin
Valrubicin
Epirubicin
LY-2874455
Nintedanib
Vorinostat
Doxorubicin
mean DSS
resistant
sensitive
T1 biomarker prediction
B)
Wilcoxon, p = 1
Anthracyclines
Chemotherapy
Azacitidine
Docetaxel
Neratinib
mean DSS
resistant
sensitive
T1 biomarker prediction
C)
biomarker present
yes
no
sample
T1
T2
Anthracyclines
Chemotherapy
Azacitidine
Docetaxel
Neratinib
mean DSS
resistant
sensitive
Biomarker prediction
D)
gene harbors somatic mutation
no
yes
Wilcoxon, p = 0.82
Wilcoxon, p = 0.52
mean DSS
T1
T2
E)
gene DE compared to benign
yes
no
Wilcoxon, p = 0.62
Wilcoxon, p = 0.048
mean DSS
FC down
FC up
F)
T1 only
as T2 had no CNVs in any of the genes targeted by the DSRT
Kruskal-Wallis, p = 0.23
0.091
0.3
0.76
mean DSS
diploid
gain
high gain
gene CN status

**Supplement Figure 7. Association of gene-level molecular features with *ex vivo* drug sensitivity scores (DSS). A-B)** Mean DSS values for T1 PDCs for drugs predicted—based on genomic biomarkers (SNVs/indels or CNVs)—to confer sensitivity or resistance. Where a biomarker mapped to a drug class (e.g., anthracyclines), DSS values were averaged. Only one drug (tamoxifen) fell into the resistance-predicted category. No significant difference in DSS was observed between predicted sensitive and resistant groups (panel A: Wilcoxon p = 0.93; panel B: p = 1). **C)** Comparison of DSS values for biomarker-predicted sensitive drugs between T1 (which harbored the relevant biomarkers) and T2 (which did not). T2 did not contain any predictive biomarkers; thus, this panel compares response to the same drug set across tumor regions. DSS values for several drugs were higher in T2 than T1, despite the absence of the associated biomarkers. **D)** Relationship between gene-level somatic mutation status and mean DSS in T1 and T2. Each point represents a gene targeted by at least one drug in the DSRT panel. Gene sets are grouped by whether the gene harbors a somatic mutation (red) or not (light pink). No significant differences in DSS were detected in either tumor region (T1: Wilcoxon p = 0.82; T2: Wilcoxon p = 0.52). **E)** Mean DSS grouped by differential expression status relative to benign kidney tissue. Genes significantly downregulated (FC down) or upregulated (FC up) are compared with non-DE genes. Upregulated genes show a modest increase in DSS (Wilcoxon p = 0.048), whereas no significant difference was detected for downregulated genes (Wilcoxon p = 0.62). **F)** Association between gene-level copy-number status and mean DSS in T1. T2 is not included because none of the DSRT-targeted genes harbored CNVs in T2. Genes are grouped as diploid, gain (at least one low-level copy-number increase), or high gain (at least one high-level amplification). Differences in DSS between groups were not statistically significant (Kruskal–Wallis p = 0.23; pairwise p-values shown above).
